# The Histone H1-like protein AlgP facilitates even spacing of polyphosphate granules in *Pseudomonas aeruginosa*

**DOI:** 10.1101/2021.08.24.457604

**Authors:** Ravi Chawla, Steven Klupt, Vadim Patsalo, James R Williamson, Lisa R Racki

## Abstract

Synthesis of polyphosphate (polyP) is an ancient and universal stress and starvation response in bacteria. In many bacteria, polyP chains come together to form granular superstructures within cells. Some species appear to regulate polyP granule subcellular organization. Despite the critical role of polyP in starvation fitness, the composition of these structures, mechanism(s) underpinning their organization, and functional significance of such organization are poorly understood. We previously determined that granules become transiently evenly spaced on the cell’s long axis during nitrogen starvation in the opportunistic human pathogen *Pseudomonas aeruginosa*. Here, we developed a granule-enrichment protocol to screen for polyP granule-localizing proteins. We identified AlgP as a protein that associates with polyP granules. We further discovered that AlgP is required for the even spacing of polyP granules. AlgP is a DNA-binding protein with a 154 amino acid C-terminal domain enriched in ‘KPAA’ repeats and variants of this repeat, with an overall sequence composition similar to the C-terminal tail of eukaryotic histone H1. Granule size, number, and spacing are significantly perturbed in the absence of AlgP, or when AlgP is truncated to remove the C-terminus. The Δ*algP* and *algP*ΔCTD mutants having fewer, larger granules. We speculate that AlgP may contribute to spacing by tethering polyP granules to the chromosome, thereby inhibiting fusion with neighboring granules. Our discovery that AlgP facilitates granule spacing allows us for the first time to directly uncouple granule biogenesis from even spacing, and will inform future efforts to explore the functional significance of granule organization on fitness during starvation.

**IMPORTANCE:** The mechanisms underpinning polyP’s pleiotropic effects on bacterial starvation physiology remain elusive. This simple polyanion’s lack of protein binding specificity has impeded validation of *bona fide* polyP-binding proteins. However, polyP forms granule superstructures with spatial specificity. Our granule enrichment protocol identified a polyP granule-associated protein in *Pseudomonas aeruginosa*, AlgP. AlgP was originally reported as regulator of alginate, an extracellular polysaccharide important in biofilm formation, including in cystic fibrosis (CF) chronic infections. AlgP’s putative role in alginate biosynthesis has recently been called into question. We establish a distinct, previously unknown function for AlgP in modulating the subcellular organization of polyP, another polymer important for pathogenesis. In CF clinical isolates, the C-terminal repeat domain of AlgP is a hotspot for genetic rearrangements. Our finding that the C-terminus of AlgP is required for granule organization lays the groundwork for exploring the functional significance of these mutations in the evolutionary trajectory of chronic infections.

## INTRODUCTION

In response to diverse starvation cues, bacteria almost universally do a curious thing: They spend ATP to make an inorganic polymer of phosphoryl groups, polyphosphate (polyP). The ability to make polyP has been shown to be important for fitness during starvation in evolutionarily disparate bacterial species, but the molecular mechanism(s) underpinning these effects has remained mysterious(1, 2). A significant challenge to understanding how polyP exerts its pleiotropic effects on physiology has been the small number of validated polyP-binding proteins identified in bacteria. Aside from the polyP biosynthetic machinery (kinase families Ppk1 and Ppk2) and a phosphatase (Ppx), only a few proteins are known to functionally interact with polyP in bacteria(1, 3). These include enzymes that have been shown *in vitro* to be able to use polyP as a substrate for phosphorylation reactions, such as NAD kinases and glucokinases(1, 3). In addition, several proteins containing a positively-charged **<u>C</u>**onserved **<u>H</u>**istidine **<u>α</u>**-helical **<u>D</u>**omain (CHAD), have been shown by microscopy to localize to polyP granules(4). It may be that the specificity of binding to polyP is mediated by cooperative assembly of multicomponent protein complexes, making it difficult to understand polyP binding through binary interactions. Because polyP forms granular superstructures in many bacteria, co-localization studies can serve as a powerful method to identify true polyP-interacting proteins.

We and others have observed that polyP granules are organized within cells, raising the possibility that the subcellular localization of these structures is a regulated process(5–8). An important motivation for characterizing the polyP granule proteome is therefore to identify not just polyP-utilizing proteins, but those that specifically contribute to the formation and organization of these structures. In *P. aeruginosa*, polyP granules become transiently evenly spaced on the long axis of the cell during nitrogen starvation. In addition to the even-spacing phenomenon, a notable structural feature of polyP granules is that they form in the nucleoid region of the cell, where the chromosome resides. PolyP granules appear to affect chromosome function, as *P. aeruginosa* cells that cannot make polyP are deficient in exiting the cell cycle during starvation, and these cells also have an elevated SOS DNA damage response(9). PolyP also affects cell cycle progression in *Caulobacter crescentus*(10). Moreover, in *C. crescentus*, it has been shown that disrupting cell cycle progression can in turn alter granule organization (*C. crescentus* cells typically have 1-2 granules, which form and localize to the quarter-cell position during cell cycle progression)(5). Taken together, these observations suggest that polyP granules may be structurally and functionally linked to the chromosome. How and why polyP is connected to bacterial chromatin remains to be explored, but the proteome of polyP granules may provide clues.

In this study, our goal was to develop a polyP granule enrichment protocol in *P. aeruginosa* strain UCBPP_PA14 (herein referred to as just “PA14”) as a screening tool to characterize the polyP granule associated proteome. We then validated our protocol by confirming that one of our candidate proteins from this screen, AlgP, indeed localizes to granules. AlgP is a histone H1-like putative DNA binding protein that has a predicted intrinsically disordered C-terminal domain containing tandem repeats of KPAA residues(11, 12). We found that the C-terminus of AlgP is required for granule spacing but that AlgP does not play a role in efficient cell cycle exit during nitrogen starvation. Our proteomic screen represents the first step in determining the interactome of polyP in the opportunistic human pathogen *P. aeruginosa*, and hopefully other bacteria in the future.

## RESULTS

### PolyP granule enrichment and proteomic analysis

We developed a protocol for enrichment of polyP granules and analysis of granule-associated proteins. We had two objectives when choosing a culture condition for granule isolation: (1) a condition in which polyP granules are evenly spaced, because we are interested in identifying protein factors that may contribute to spatial positioning of granules, and (2) a condition where polyP granules are abundant, and the presence of polyhydroxyalkanoate (PHA) granules is minimized, to avoid cross-contamination by proteins associated with PHA granules. Three hours of nitrogen starvation in minimal medium satisfied both of these criteria, as shown in our previous work(9). Briefly, we fractionated a lysate of WT and a ΔpolyP strain (a quadruple knockout of *ppk1, ppk2A, ppk2B*, and *ppk2C*) by ultracentrifugation in a self-forming Percoll gradient(13)(Fig 1A). We observed a brownish-yellow pellet at the bottom of the WT gradient, but not the ΔpolyP gradient (Fig 1B). We extracted proteins from the pellet and whole lysate of WT and subjected them to label-free proteomic analysis (see Materials and Methods for details).

**Figure 1.**
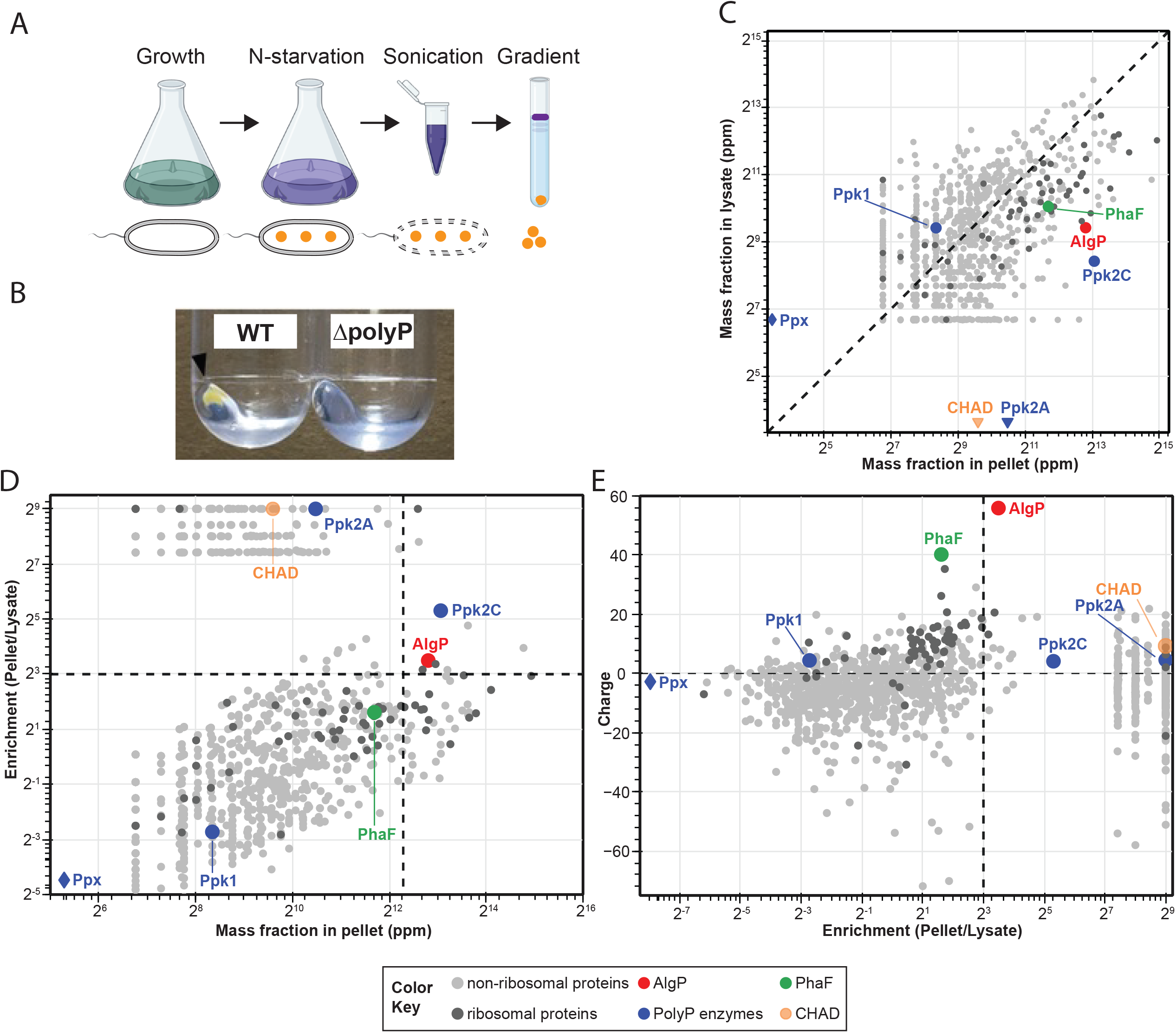
Workflow for isolation of polyphosphate granules from starved *P. aeruginosa* cells and proteomic analysis. **(A)** The workflow of granule isolation as described in the Methods section. **(B)** The ultracentrifugation of a lysate from wild type *Pseudomonas aeruginosa PA14* cells in a Percoll gradient solution generates a “pellet” fraction, which is not seen with a lysate from ΔpolyP strain. The bottom panel shows a zoomed-in view of the Percoll-encased pellet after discarding the Percoll gradient solution. **(C-E):** Mass spectrometry analysis of the pellet fractions from three independent experiments. **(C)** Absolute mass fraction of proteins in the “pellet” and “lysate”, obtained from spectral counting (see Methods and Fig S1) is shown in parts per million. The dashed diagonal corresponds to a line of equal abundance in the pellet and lysate fraction. The data points below the diagonal line correspond to proteins in higher abundance in the pellet fraction than in the lysate; diamond and inverted triangle symbol marks the proteins identified in “lysate-only” and “pellet-only” fractions. (**D)** The enrichment of proteins in the pellet plotted against the abundance of proteins in the pellet. The cut-off of lines for enrichment are 8, (2^3^, log_2_ fold-change =3) and abundance of 5000ppm. The highly abundant and enriched proteins in the pellets are, therefore, located in the upper right quadrant. **(E)** The calculated charge at neutral pH for proteins identified in the proteomics plotted against the fold enrichment (negative charge cut-off for plotting is -75; refer to Fig S1c for the full plot). The horizontal dashed line divides the data into positively and negatively charged proteins. For panels C-E: averages of proteins identified in the mass spectrometric analysis from three independent experiments (light gray circles); ribosomal-binding proteins (dark gray circles); AlgP(red circles); polyP kinases(blue circles); polyhydroxyalkanoate granule-associated protein PhaF(green circles); and CHAD proteins (orange circles).

We identified approximately 1200 proteins in the pellet fraction. Average protein abundances for the pellet and lysate from three independent experiments are shown in Figure 1C. As an initial verification for extraction methodology, we sought to confirm enrichment of known polyP associated proteins, including the previously mentioned polyP kinases (Ppk1, Ppk2A, Ppk2B, and Ppk2C), exopolyphosphatase (Ppx), and two CHAD-containing proteins (PA14_13320 and PA14_26940). Ppk1 was more abundant in the lysate than in the pellet and Ppk2B was not detected in our screen. Ppk2C was enriched and highly abundant in the pellet fraction. Ppk2A was detected in the pellet fraction in one of three replicates with low abundance. Ppx was present in the lysate fraction only. One of the CHAD-containing proteins, PA14_26940, was consistently detected in the pellet fraction but PA14_13320 was not. We note that some ribosome-associated proteins were more abundant and enriched in the pellet than lysate. However, we excluded this class from subsequent analysis, as they are known to easily extract under the salt conditions used for our screen(14).

We hypothesized that our granule proteomic screen would identify two broad but non-exclusive classes of proteins: (1) structural proteins that contribute to polyP granule formation, and (2) client proteins that use polyP for another function, such as kinases that use polyP for phosphorylation reactions. PolyP has one of the highest charge densities of any macromolecule in the cell(15). Hence, we decided to focus initially on the first class, which we reasoned would have two key characteristics: high enrichment and positive charge. A plot of enrichment, defined as the ratio of abundance of protein in pellet to the lysate, versus the abundance is shown in Fig 1D. Ppk2C was one of the nine highly enriched and abundant enriched protein candidates identified in this step. A plot of net protein charge at pH 7 (obtained from the Pseudomonas Genome Database (16)) versus protein enrichment in the polyP pellet is shown in Figure 1E, revealing one protein, AlgP, that distinctly stood out due to its net charge of +55. In contrast, PhaF, which is a polyhydroxyalkanoate (PHA) granule associated protein with a net positive charge +35 was abundant, but not enriched per our screening criteria. AlgP and PhaF share a highly homologous C-terminal domain enriched in closely-spaced lysine residues (see Discussion), and their abundance is therefore likely an underestimate due to the trypsin cleavage treatment for mass spectrometry. Based on these abundance, enrichment, and physiochemical criteria, we decided to focus on AlgP for subsequent analysis.

### AlgP localizes to polyP granules

We used fluorescence microscopy to assess whether AlgP is enriched in polyP granules, as our proteomic analysis suggests. We replaced the endogenous copy of AlgP in the genome with N and C-terminally tagged fluorescent chimeras of AlgP with mApple. AlgP-mApple forms small irregular puncta and patches in exponentially growing cells (Fig 2A, top left). After 3h of nitrogen starvation, AlgP-mApple forms discrete evenly spaced puncta on the long axis of the cell (Fig 2A, bottom left). Total fluorescence of AlgP-mApple revealed that AlgP is expressed during exponential growth and increases in abundance during starvation (Fig 2B).

**Figure 2:**
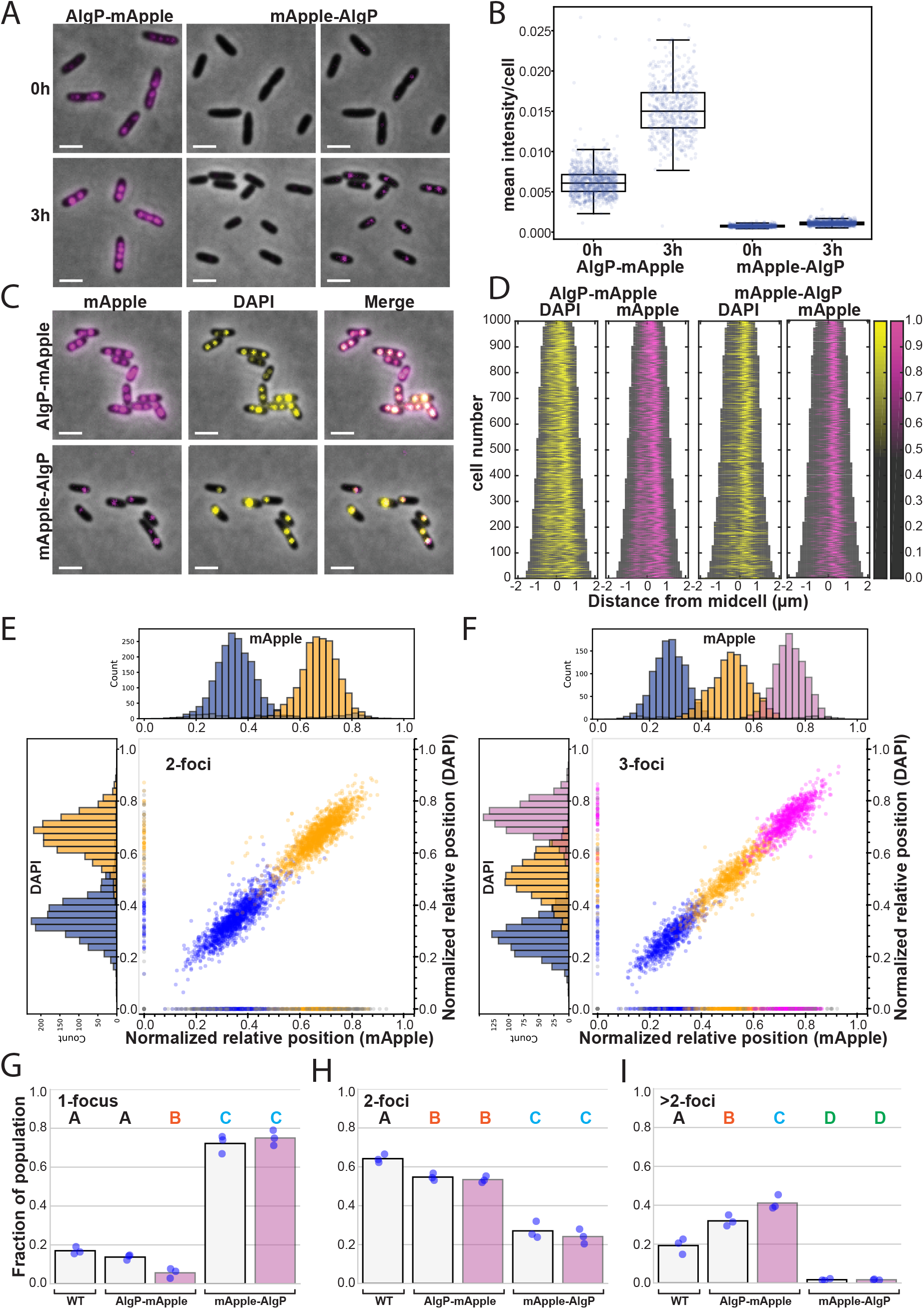
Subcellular localization of AlgP. (A) Representative images of localization of AlgP-mApple and mApple-AlgP translational fusion proteins. The genes encoding the chimeric proteins replaced the endogenous *algP* at its native chromosomal locus. mApple fluorescence displayed in magenta. Left panels: AlgP-mApple in exponential growth (0h) and after nitrogen starvation (3h). Middle panels: mApple-AlgP displayed using the same contrast settings as for AlgP-mApple. Right panels: the same mApple-AlgP images, shown with the higher contrast settings necessary to see puncta. (B) Quantification of the average fluorescence intensity per cell (total intensity divided by the area of the cell) for AlgP-mApple and mApple-AlgP in growing (0h) and nitrogen-starved (3h) cells. (C) Localization of AlgP-mApple and mApple-AlgP relative to DAPI-stained polyP granules. mApple fluorescence is displayed in magenta, DAPI fluorescence is displayed in yellow. Note that the contrast settings in mApple channel are different for AlgP-mApple and mApple-AlgP because of different brightness. They reflect the same settings used in the left panels of (A) for AlgP-mApple, and the right panels of (A) for mApple-AlgP. (D) Demographs displaying the fluorescence intensity of DAPI and mApple on the long axis of the cell in 3h nitrogen-starved cells. (E) Spatial organization of 2-foci cells. Histograms display the normalized relative position on the long axis of the cell of the first foci in blue and second foci in orange. The horizontal histogram (top) represents mApple foci, and the vertical (left) histogram represents DAPI-labeled foci. Dots in the scatter plot show the correlation in spatial position of mApple and DAPI foci. Grey dots represent foci found in one channel but not in the other, these are also displayed as grey in the histograms. (F) Spatial organization of 3-foci cells. As in (E), but the third foci is depicted in magenta. (G) Quantification of the fraction of 1-foci cells in the population in both the DAPI and mApple channel. Note that 0-puncta cells are not included in these calculations, because it is impossible to distinguish between cells that have no granules, and those in which DAPI did not efficiently label the cell. The fraction of 1-foci cells therefore represents the ratio of 1-foci cells to all cells with 1 or more foci. Each point represents an independent replicate/day of experiment. Dots represent results from independent experiments on different days, bar shows mean. Variance analyzed using a one-way ANOVA. Significant differences between strains at the same time point are marked with different uppercase letters based on a post hoc Tukey test (H) As in (G) but for 2-foci cells. (I) As in (G) but for cells with more than 2 foci. Scale bar: 2µm.

We then used DAPI staining to compare the localization of these puncta with polyP granules, and qualitatively observe clear co-localization (Fig 2C, top panels). Demographs of the relative position of DAPI and AlgP-mApple-labeled foci on the long axis of the cell reveal a similar banding pattern between the two fluorescent channels (Fig 2D). We compared the number and positioning of DAPI-labeled polyP granules in WT versus AlgP-mApple cells, for all foci-containing cells, to determine if the chimeric fusion interferes with granule biogenesis. To determine whether the even spacing granule phenotype we have previously observed is preserved in AlgP-mApple-labeled cells, and to quantify co-localization of DAPI and AlgP-mApple labeled foci, we examined the normalized relative position of foci on the long axis of the cell in both channels, for both 2-foci and 3-foci cells (Fig 2E, F). We observed even spacing in both channels, and the position of individual foci in the two channels was positively correlated in the scatter plot. We also compared the number of foci per cell in cells expressing AlgP-mApple with that of WT, again for all foci-containing cells. We observed a similar fraction of cells with 1 DAPI-labeled foci in WT and AlgP-mApple cells (Fig 2G), but slightly more 2 DAPI-labeled foci in WT than AlgP-mApple cells, and slightly fewer >2 DAPI-labeled foci in WT than AlgP-mApple cells (Fig 2H, I). The fraction of mApple-labeled foci per cell was similar to DAPI-labeled foci for 2-foci cells, though we observed slightly fewer 1 mApple-labeled foci than 1 DAPI foci per cell, and slightly more >2 mApple-labeled foci than >2 DAPI labeled foci (Fig 2G-I, summarized in Table S3a).

The mApple-AlgP chimera resulted in a different behavior. The total fluorescence intensity of this protein was significantly lower both in exponential growth and during starvation, suggesting that the N-terminal tag may interfere with expression, folding, or stability of the full-length protein (Fig 2B). The localization of mApple-AlgP was different from AlgP-mApple: we still observed puncta, but significantly fewer than we observed with the C-terminal tag (Fig 2A, right panels). The mApple-AlgP protein co-localize with granules, but granule number and organization are significantly perturbed (Fig 2C, bottom panels, D, G). Together these observations suggested that AlgP might play a role in the subcellular positioning of polyP granules.

### AlgP affects granule maturation and spatial organization

AlgP is a 352 amino acid (aa) protein with a 105aa N-terminal domain predicted to be α-helical (Phyre2), and a 195aa predicted intrinsically disordered C-terminal domain containing a 154aa region of tandem repeats of KPAA(17). There are 25 perfect KPAA repeats interspersed with variants, including 7 KPVA, 4 KTAAA, one KPAV, and two alanine spacers. Two KPAA repeats fall outside of this 154aa contiguous region (Fig 3A and S2). To determine if AlgP is required for polyP granule formation and/or organization, we constructed a clean deletion (Δ*algP*). Granule number and spatial organization is perturbed in Δ*algP* cells (Fig 3B, C). We performed complementation analysis by inserting a copy of *algP* under its native promoter at the *att*Tn7 locus, and observed partial rescue of granule number and organization (Fig S3). Given the limits of light microscopy to detect small granules and to accurately measure granule size, we used TEM as an orthogonal method to characterize the granule phenotype in Δ*algP* cells early in starvation (1.5h) and at 3h. We observed fewer, larger granules for Δ*algP* cells at both timepoints (Fig 4A-C). While granules were larger in Δ*algP* cells than WT cells at 1.5h, we saw a larger subpopulation of cells without granules at this timepoint in Δ*algP* cells (26% of ΔalgP cells versus 2% of WT cells). Even including these ‘zero-volume’ cells in the distribution, the average volume of granules for Δ*algP* cells was still larger than for WT cells. Indeed, when we look at the volume of the largest granule per cell, this trend towards larger granules becomes very apparent, (Fig 4D). While the distribution of granular volume is different in Δ*algP* cells, with fewer, larger granules, the loss of AlgP does not appear to lead to a net decrease in the total volume of polyP granules per cell. We see a broader distribution of total granular volume per cell Δ*algP* cells than in WT cells (Fig 4E). Summary statistics for TEM quantification are provided in Table S3b.

**Figure 3.**
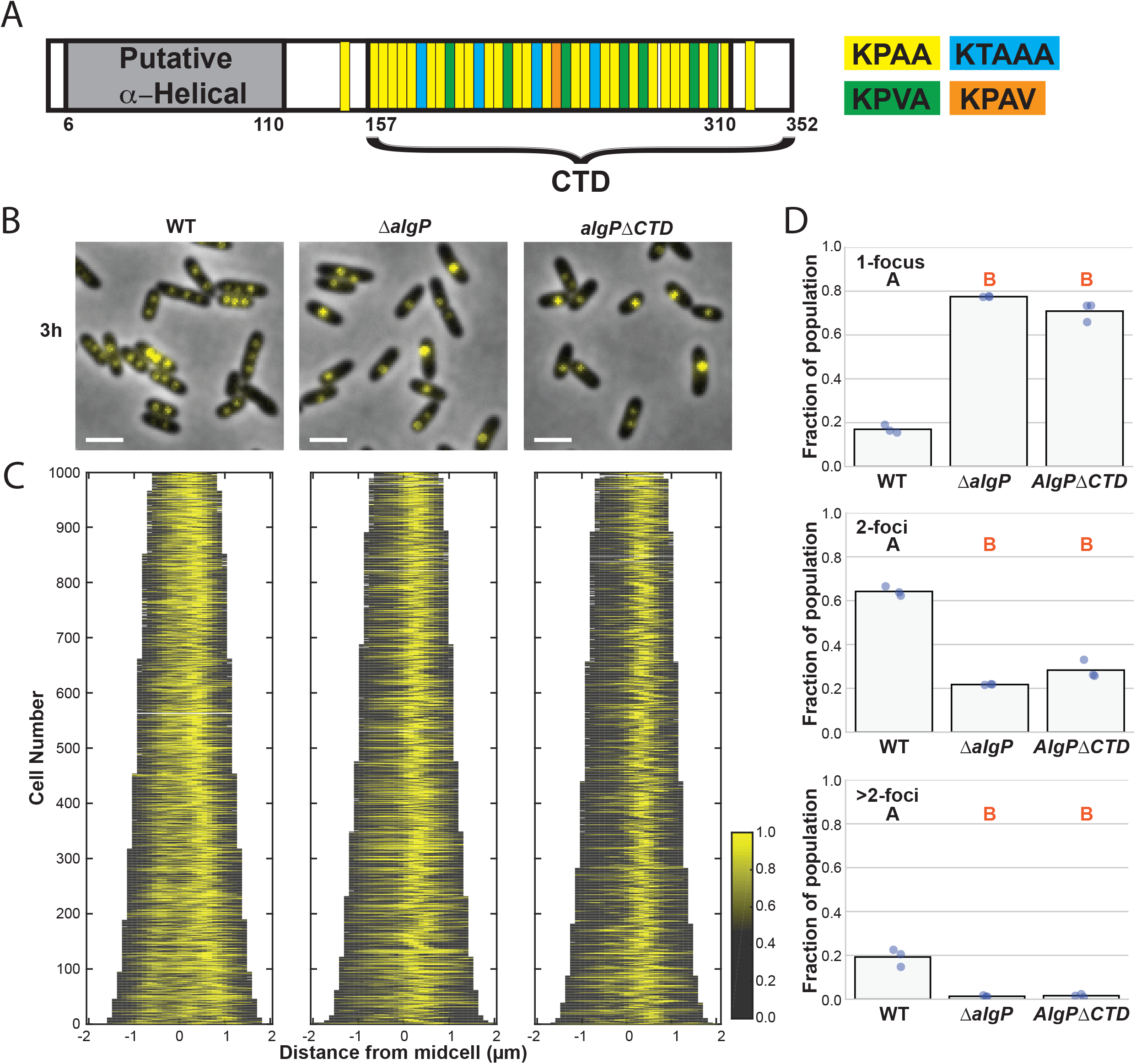
Effect of AlgP and C-terminus of AlgP on localization of polyP granules during nitrogen starvation. (A) AlgP protein domain organization. AlgP contains an N-erminal 105aa putative α-helical domain, predicted by Phyre2. The C-terminus contains a 195aa domain predicted to be intrinsically disordered, with a 154 region of tandem repeats of KPAA. There are 25 perfect KPAA repeats interspersed with variants, including 7 KPVA, 4 KTAAA, one KPAV, and two alanine spacers. Two KPAA repeats that fall outside of this 154aa contiguous region. (B) Representative images of DAPI-stained cells. (C) Demographs displaying the fluorescence intensity of DAPI on the long axis of the cell in 3h nitrogen-starved cells. (D) Quantification of the fraction of DAPI-labeled cells in the population with 1, 2, and >2 foci per cell. Each point represents an independent replicate/day of experiment. Note that 0-puncta cells are not included in these calculations, because it is impossible to distinguish between cells that have no granules, and those where DAPI did not efficiently label the cell. The fraction of 1-foci cells therefore represents the ratio of 1-foci cells to all cells with 1 or more foci. Dots represent results from independent experiments/days, the bar indicates the mean. Variance analyzed using a one-way ANOVA. Significant differences between strains at the same time point are marked with different uppercase letters based on a post hoc Tukey test. Scale bar: 2µm.

**Figure 4.**
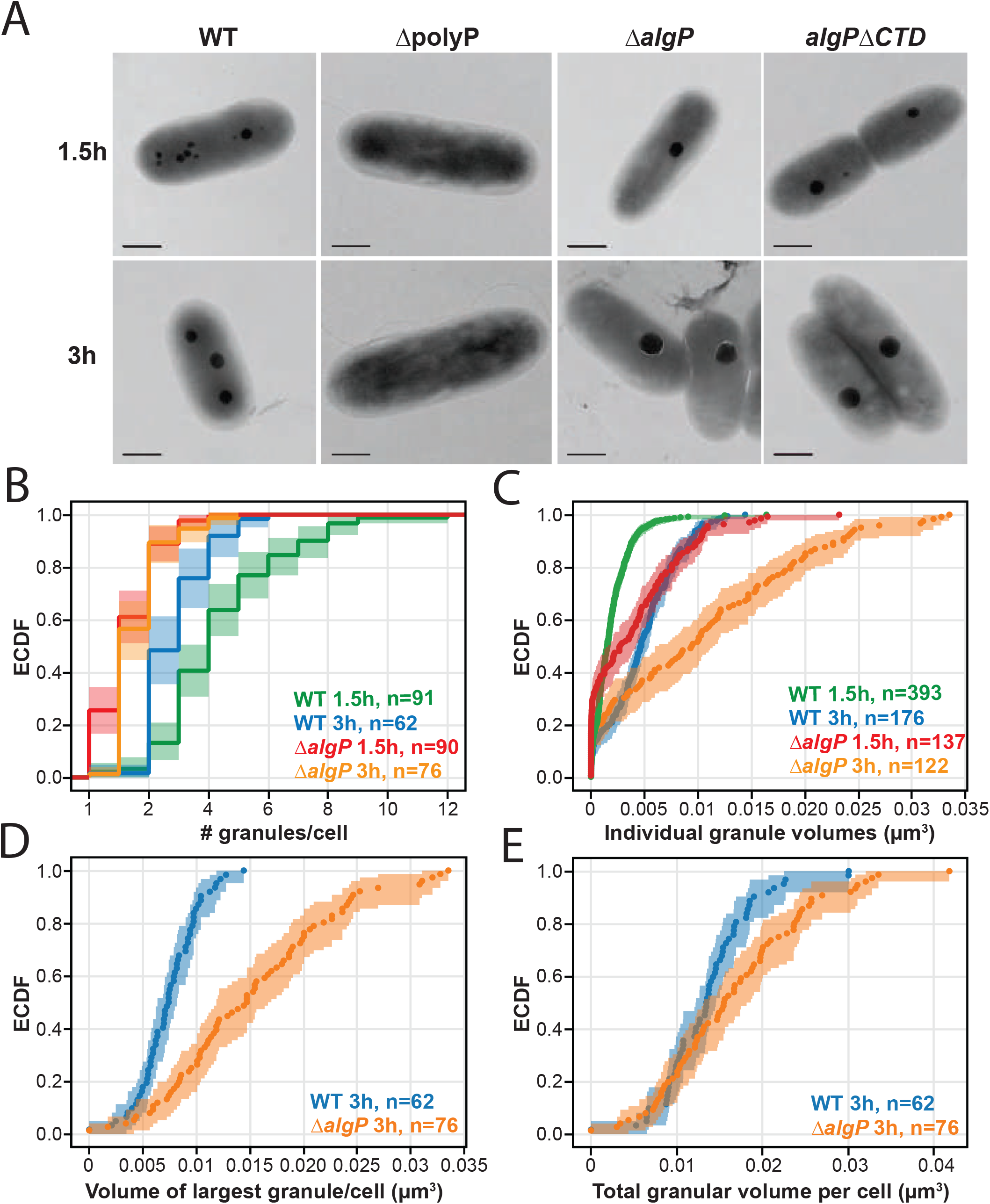
Effect of AlgP and C-terminus of AlgP on number and size of polyP granules during nitrogen starvation. For the empirical cumulative distribution functions (ECDFs), the shading represents confidence intervals for the ECDF acquired by bootstrapping. (A) Representative TEM images of cells 1.5 and 3h after nitrogen starvation. (B) Distribution of the number of granules per cell for WT and Δ*algP* cells at 1.5 and 3 hours into nitrogen starvation. (C) Distribution of volumes of all individual granules at 1.5 and 3 hours into nitrogen starvation. (D) Distribution of volumes of the largest granule per cell after 3 hours of nitrogen starvation. (E) Distribution of total granular volume per cell (sum of volumes of all granules within a cell) after 3 hours of nitrogen starvation. Scale bar: 0.5µm.

We note that for the TEM analysis we observe a subpopulation of cells with many small ‘satellite’ granules. We previously observed these small granules of 59 ± 20 nm nanometers in diameter, or smaller, appear only in a fraction of cells(9). We did not quantify these ‘satellite’ granules in our analysis, but we note the fraction of cells that contain them: in 3h nitrogen-starved cells, we observe these in 24% of WT cells and 26% of Δ*algP* cells. We show examples of cells with ‘satellite’ granules in Fig S4A.

### Deletion of the KPAA repeat domain results in loss of granule positioning

The C-terminal 154-amino acid KPAA repeat domain of AlgP is highly homologous to the C-terminus of PhaF. In PhaF, this domain is thought to bind DNA and tether polyhydroxyalkanoate (PHA) granules to the nucleoid and thus contribute to their even distribution within cells. To determine if the KPAA-repeat domain of AlgP plays a role in granule spacing, we generated an AlgP truncation mutant where we removed the last 195 C-terminal amino acids (residues 157-352, including the 154aa repeat region, Fig 3A, S2), In this *algP*ΔCTD strain, we observe a similar decrease in granule number per cell as with Δ*algP* cells (Fig 3A-C). Qualitatively we also observe this decrease in granule number by TEM (Fig 4A, right panels) though we have not quantified this effect in the *algP*ΔCTD strain for the TEM data.

### AlgP localization depends on polyP kinases

In the absence of starvation, we noticed that AlgP-mApple is not diffuse, but already forms puncta (Fig 2A). These puncta might form due to interactions with the chromosome, or with a specific polyP kinase, or nascent polyP granules. Indeed at 0h we observed small putative nascent granules by TEM, though their presence in growing cells is variable (examples shown in Fig S4B). To determine whether AlgP localization is tied to the polyP polymer itself or instead to the polyp kinases, we expressed our AlgP-mApple fusion in various *P. aeruginosa* strains lacking some or all of the kinases. A mutant lacking all four polyP kinases lacks these puncta, suggesting that the localization depends on polyP and/or specific polyP kinases (Fig 5A). We note that in ΔpolyP cells, we occasionally observe cells with a single DAPI stained puncta (<10% of cells), with which AlgP-mApple co-localizes. We do not believe these puncta are polyP granules because of their small size and number per cell, and because we do not see them by TEM (Fig 4A). We can only speculate about what they might represent, acknowledging that dyes can act non-specifically and DAPI is known to undergo a red emission shift when bound to RNA as well as other biomolecules(18). We next sought to determine whether the AlgP-mApple puncta we observe depend on the polyP polymer, or one of the polyP kinases specifically. We previously showed that either Ppk1 or Ppk2A is sufficient for granule biogenesis(9). In a Δ*ppk2a*Δ*ppk2b*Δ*ppk2c* triple mutant, we observe puncta formation, indicating that Ppk1 alone is sufficient for AlgP-mApple puncta formation (Fig 5A, 3^rd^ column). Similarly, in a Δ*ppk1*Δ*ppk2b*Δ*ppk2c* triple mutant, we observe puncta formation, indicating that Ppk2A alone is also sufficient (Fig 5A, right panels).

**Figure 5.**
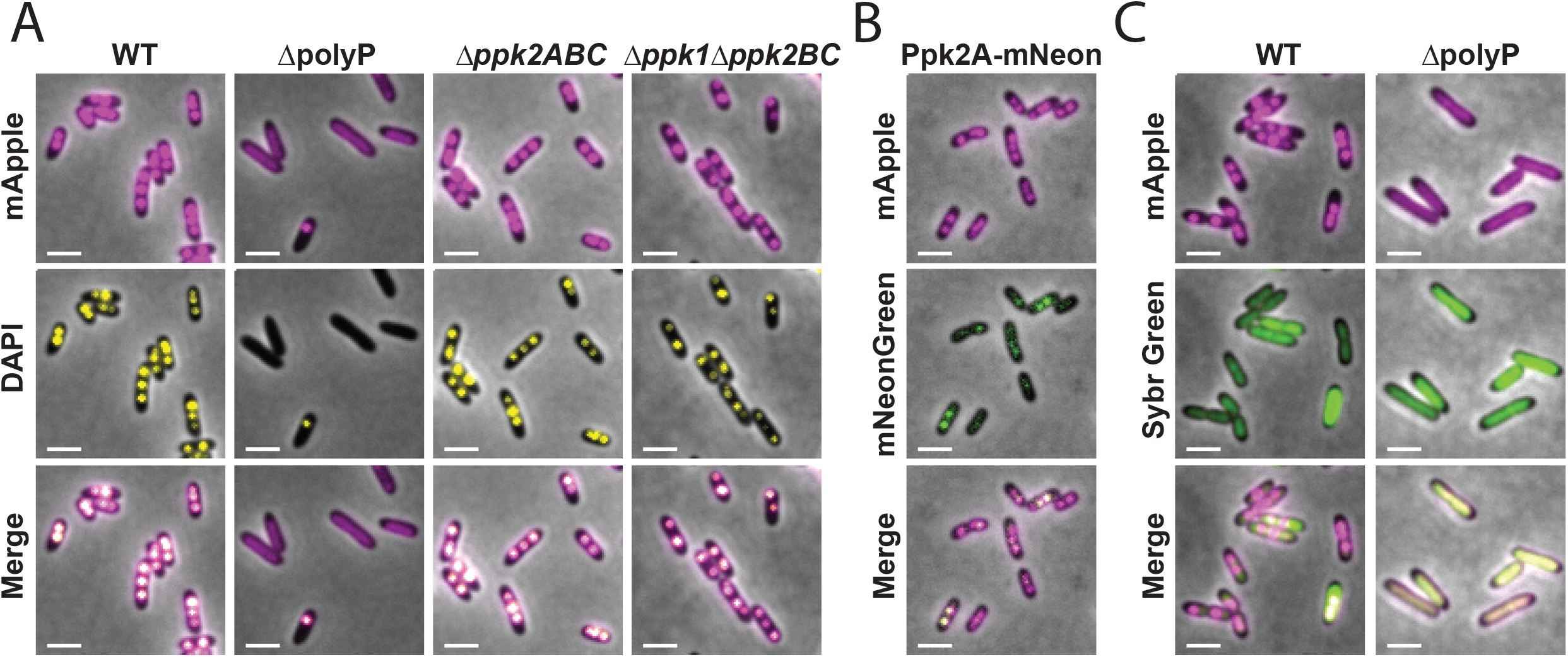
Effects of polyP kinases on AlgP and polyP granule localization after 3h of nitrogen starvation. (A) Localization of AlgP and DAPI-labeled puncta in the absence of all polyP kinases (ΔpolyP), when only Ppk1 is present (3^rd^ column), and when only Ppk2A is present (4^th^ column). (B) Localization of AlgP-mApple relative to Ppk2A-mNeonGreen. (C) Localization of AlgP relative to the nucleoid in WT and ΔpolyP cells. The nucleoid was visualized using Sybr Green labeling (2^nd^ row). Scale bar: 2µm.

We then looked at co-localization of AlgP-mApple with Ppk2A. We previously showed that Ppk2A-mCherry can co-localize with DAPI-labeled polyP granules(9). We qualitatively observe co-localization of Ppk2A-mNeonGreen with AlgP-mApple, although the Ppk2A-mNeonGreen and Ppk2A-mCherry signals are low when these proteins are expressed from the native *ppk2A* promoter and not every DAPI-stained or AlgP-mApple-labeled granule has visible Ppk2A fluorescence (Fig 5B). We conclude that AlgP-mApple is a brighter and more consistent marker of granules than Ppk2A-mNeonGreen.

Finally, we noticed that the AlgP-mApple fluorescence in ΔpolyP cells was not completely diffuse throughout the cytoplasm (Fig 5A, 5C). Instead, we observed apparent co-localization of AlgP-mApple with the nucleoid region of the cell, as visualized by Sybr Green staining, suggesting that in ΔpolyP cells, AlgP may bind nonspecifically to DNA (Fig 5C). Together these data suggest that AlgP-mApple puncta formation depends not on a specific polyP kinase, but rather the presence of small amounts of polyP in growing cells, and that AlgP may bind to the chromosome more broadly in the absence of polyP.

### AlgP is not required for efficient cell cycle exit

We previously showed that polyP is required for efficient cell cycle exit in response to nitrogen starvation(9). We also previously showed that transient even spacing of granules on the long axis of the cell coincides with the period of cell cycle exit, between 3 and 6 hours into nitrogen starvation(9). We therefore wondered whether AlgP, which is required for granule organization, might also affect efficient cell cycle exit. To characterize the effect of AlgP on cell cycle exit, we used fluorescent markers to label the chromosomal origin of replication and open DNA replication forks, as we have done previously. Briefly, we labeled open DNA replication forks with a fluorescent chimera of single stranded DNA binding protein (SSB-mCherry), and we used the heterologous chimeric chromosome partitioning protein GFP-ParB^pMT1^ and its cognate DNA binding sequence *parS*^pMT1^ to label the chromosomal origins. In exponential growth, most WT and ΔpolyP cells are copying their chromosomes, with more than one origin/cell and an open DNA replication fork (Fig 6B,C, summarized in Table S3c). While we observe a similar fraction of cells with open forks in exponentially growing Δ*algP* cells, the fraction of cells with more than one origin per cell is significantly lower. However, the bulk growth rates of the WT and Δ*algP* are similar (Figure S5), suggesting that the difference we observe with the fluorescent tags does not reflect an underlying replication defect. At six hours after shifting exponentially growing cells to nitrogen-limited MOPS minimal media, 10±6% of WT and 13±5% of Δ*algP* cells have more than one origin per cell, and 3±2% of WT and 2±2% of Δ*algP* cells have open DNA replication forks (Fig 6B). In contrast, as we previously have shown, ΔpolyP cells have a defect in efficient exit, as 87±4% still had more than one origin/cell and 35±3% had open forks after 6 hours of nitrogen starvation (Fig 6B,C, summarized in Table S3c). These data suggest that AlgP does not play a role in efficient cell cycle exit during nitrogen starvation.

**Figure 6.**
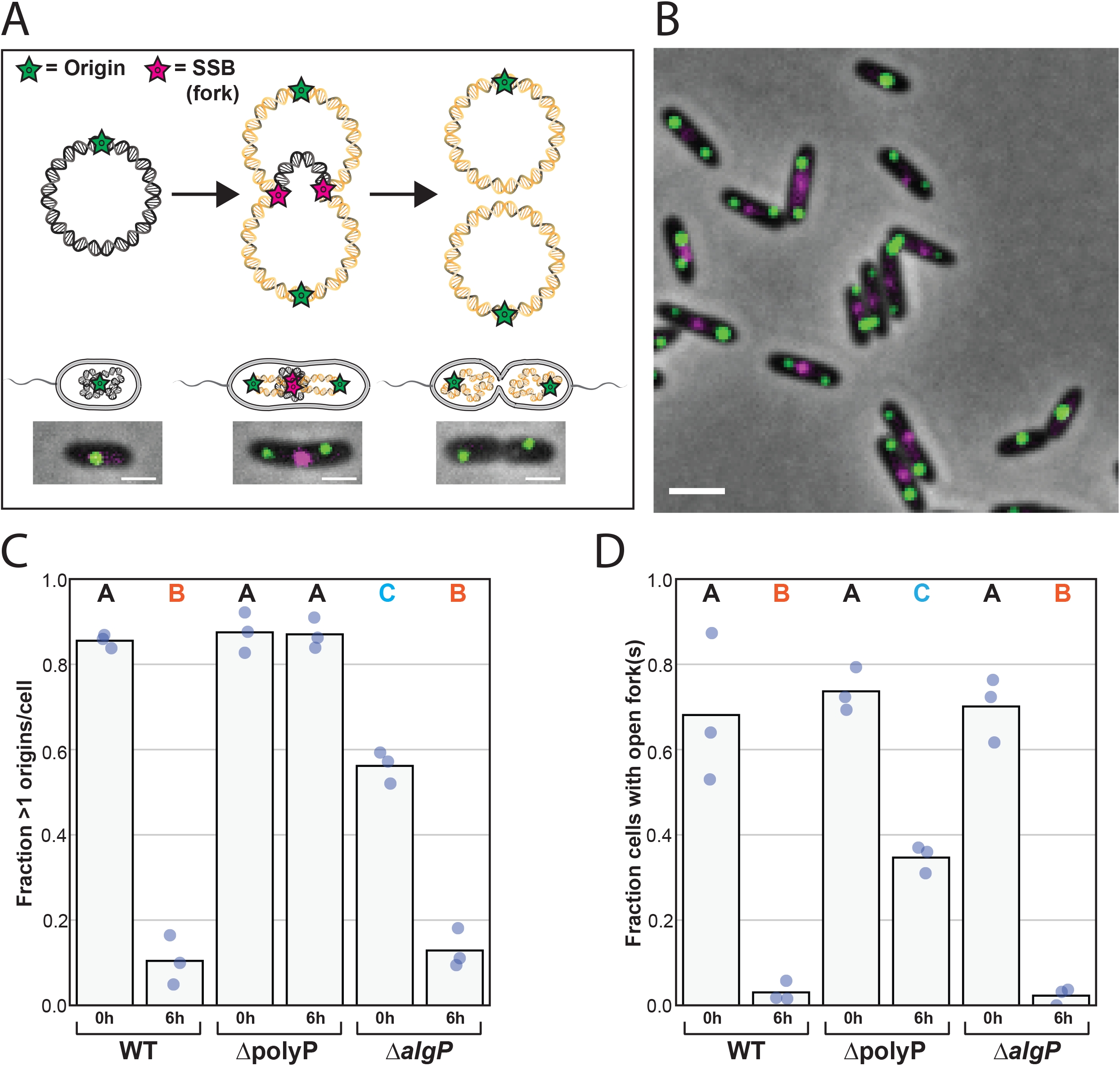
Effects of AlgP on cell cycle exit during nitrogen starvation. (A) Representative image of WT exponentially-growing cells with chromosomal origins (green) and open DNA replication forks (red). Origins are labeled using GFP-ParB^pMT1^ and a single copy of its cognate DNA binding site, *parS*^*pMT1*^ DNA sequence at the *attB* locus, 19.5 kb from the origin. Forks are labeled using a fluorescent chimera of single stranded DNA-binding protein (SSB-mCherry). We used a second copy of the SSB promoter to drive both SSB-mCherry and GFP-ParB^pMT1^ expressed from the same operon, and also integrated at the *attB* locus. (B) Fraction of cells with more than one chromosomal origin per cell in growing (0h) or 6h nitrogen starved (6h) cells. Each point represents an independent replicate/day of experiment. (C) Fraction of cells with at least one open DNA replication fork in growing (0h) or 6h nitrogen-starved (6h) cells. Each point represents an independent replicate/day of experiment. In (B) and (C), dots represent results from independent experiments/days, and the bar indicates the mean. Variance was analyzed using a one-way ANOVA. Significant differences between strains at the same time point are marked with different uppercase letters based on a post hoc Tukey test. Scale bar: 2µm.

## DISCUSSION

We and others have observed spatial organization of polyP granules in bacteria, and further hypothesized that polyP granules may have an organizational role(5, 19–22). Our granule enrichment protocol led us to identify and validate the protein AlgP as a *bona fide* polyP associated protein. We have also shown that AlgP has a structural role in granule formation: cells lacking AlgP or the C-terminal domain of AlgP lack evenly spaced granules. How AlgP promotes even spacing of granules remains to be determined, but the amino acid sequence of its C-terminus, along with previous studies of AlgP and related proteins, provide exciting clues.

### Proteomic Screen: Proof of principle and next steps

PolyP has been implicated in diverse roles in bacterial cell physiology, but the identity and function of the proteins that interact with this structurally simple biomolecule *in vivo* have remained elusive. Previous studies have reported isolation of a polyP pellet obtained from centrifugation of lysates in other bacteria including *Desulfovirbiro gigas, Corynebacterium glutamicum, and Ralstonia eutropha*(23–25). In *R. eutropha*, these methods led to enrichment of known polyP kinases, confirmed by fluorescence microscopy. To the best of our knowledge, the enrichment and characterization of polyP granule-associated proteins has not been reported in *P. aeruginosa*.

Our granule enrichment protocol identified AlgP as a polyP-granule associated protein in *P. aeruginosa* PA14. Here, the abundance and enrichment characteristics of two known classes of polyP-associated proteins (polyP kinases and conserved histidine α-helical domain (CHAD) motif containing proteins) are worth noting. *P. aeruginosa* has one known polyP phosphatase (ppx) and two classes of polyP kinases: one Ppk1 and three Ppk2-type kinases (Ppk2a, Ppk2b and Ppk2C). In our screen Ppk1 and Ppx were consistently depleted in the pellet fraction (Fig 1C). Although Ppk2A was enriched in the pellet fraction at only low abundance and was detected in only one of three independent experiments, Ppk2A-mNeonGreen colocalizes with polyP granule (Fig 5C). Hence it is possible that Ppk2A weakly or transiently associated with polyP granules and that our granule isolation protocol might have disrupted this association. Ppk2B was not detected in our screen but Ppk2C was consistently and significantly enriched and abundant. Differential association of polyP kinases and phosphosphotases with polyP granule has previously been observed in betaproteobacterium *Rastlonia eutropha* H16: Ppk1A, Ppk2C, Ppk2D, and Ppk2E localize with the polyP granule, whereas Ppk1B, Ppk2A, Ppk2B and Ppx are not associated with granules(4, 25).

Future work will seek to determine the co-localization characteristics of other proteins identified and prioritized in our current screen. Recent work has reported that long-term nitrogen starvation in *Escherichia coli* leads to the formation of large subcellular assemblies, like H-bodies and Hfq-foci, which are a part of the RNA degradosome(26). Although these assemblies have not been reported in starved *P. aeruginosa* cells, it is plausible that polyP could nucleate and mediate pelleting of these or other unknown large bodies. The availability of AlgP as a granule marker will facilitate development of improved granule fractionation and isolation protocols.

### Beyond alginate biosynthesis: A role for AlgP during starvation

AlgP was previously identified as a transcription factor thought to promote synthesis of alginate, an extracellular polysaccharide that contributes to biofilm production and when overproduced results in a ‘mucoid’ phenotype in chronic infections which correlates with a poor prognosis(27, 28)(29). The *algP* gene is adjacent *algR* and *algQ*, known regulators of alginate biosynthesis. Recently, the putative role of AlgP in alginate biosynthesis in *P. aeruginosa* has been called into question with the observation that deletion of *algP* in two different mucoid strains of *P. aeruginosa*, did not result in a nonmucoid phenotype. In the Δ*algP* mutant, alginate production was not attenuated and the alginate biosynthetic operon was not affected(30). Our discovery that AlgP affects polyP granule localization demonstrates that this protein remains important for starvation physiology. Although our AlgP mutant does not display a defect in cell cycle exit, there are intriguing clues that it may play a role in modulating gene expression and affect anaerobic survival(30). Indeed, AlgP was identified as highly expressed in the metabolically less active regions of *P. aeruginosa* biofilms using BONCAT labeling, suggesting that starvation upregulates its expression(31). In addition to localizing to granules, suggesting a direct structural role in granule organization, it is also possible that AlgP modulates granule organization indirectly via its effects on gene expression during starvation. While AlgP may not control alginate biosynthesis, polyP and alginate, two polymers associated with fitness during starvation, may nevertheless share a regulatory link. An older study implicated the transcriptional regulator AlgQ (originally called AlgR2) in polyP biosynthesis(32). An Δ*algQ* mutant of *P. aeruginosa* has reduced levels of polyP, which can be complemented either by adding back AlgQ, or one of its putative downstream effectors, nucleotide diphosphate kinase (Ndk). The authors attribute the ability of Ndk to rescue polyP synthesis to its role in modulating nucleotide pools, particularly GTP pools. GTP is required to make the stringent response regulator (p)ppGpp, which was long thought to be required or polyP synthesis. More recently, in both *E. coli* and *P. aeruginosa*, (p)ppGpp was found not to be required for polyP synthesis(9, 33). Ndk may still influence polyP biosynthesis via its effect on nucleotide pools(9, 33). Further work is necessary to explore the possible functional relationship between alginate and polyP, both important biopolymers in starvation and pathogenesis.

### The C-terminus of AlgP: connecting polyP and bacterial chromatin

TEM and cryo-electron tomography of diverse bacterial species indicate that polyP granules tend to form in the cell’s nucleoid region(5–8). How and why these negatively charged polyanions associate has remained unclear, but their colocalization suggests there may be a functional and/or structural relationship between polyP granules and bacterial chromatin. While AlgP has not previously been implicated in polyP granule biogenesis, the protein has long been thought to associate with the nucleoid. The CTD of AlgP binds DNA, and peptides containing KPAA repeats bind DNA *in vitro*(27, 34). Molecular modeling studies suggest that the CTD may form a helix around DNA(34). A second protein in *P. aeruginosa*, PhaF, contains a highly homologous CTD to AlgP and is also thought in other Pseudomonads to bind DNA(35–37). PhaF has been shown to play a role in the organization of another class of granular structures, polyhydroxyalkanoate (PHA) granules, which serve as a cellular store of reduced carbon(35, 36). PhaF is thought to tether PHA granules to the chromosome, and in its absence PHA granules clump together(36). The N-terminus of PhaF contains an amphipathic a helix believed to interact with the hydrophobic surface of PHA granules, as well as an oligomerization domain thought to drive tetramerization(35). The N-terminus of AlgP is predicted to be a helical but its structure and function is not yet known.

In addition to the sequence homology to PhaF, AlgP has interesting similarities in terms of sequence composition to the intrinsically disordered C-terminal tail of eukaryotic histone H1, also rich in lysine, proline, and alanine, as has been previously noted(12, 27, 30). Unlike the other core histones which are thought to have evolved from archaea, the histone H1 family is thought to be of bacterial origin(38). Histone H1 in eukaryotes plays a role in chromatin condensation, and its CTD interacts with linker DNA flanking the nucleosome(39). We observed that in the absence of polyP kinases, AlgP appears to globally interact with the nucleoid. Further chromatin immunoprecipitation studies are needed to understand its sequence preference *in vivo*. While the oligomeric state of AlgP is not yet known, it is possible that the N-terminus might mediate higher order oligomers that are competent to interact with both polyP and DNA.

### The C-terminus of AlgP: mediator of phase separation?

A growing list of phase separated microcompartments have been observed in bacteria and are thought to enable cells to achieve subcellular organization and generate specific microenvironments in the absence of membrane-bound organelles(40, 41). Formation of such compartments is driven by polyanions such as RNA, and intrinsically disordered proteins such as PopZ(42–46). Recently, polyP has been shown to form liquid-like condensates with polycationic peptides and supercharged GFP (+36GFP)(47–49). Additionally, the synthetic +36GFP-polyP condensates were shown to coalesce and exhibit liquid-like properties in *Citrobacter freundii in vivo*(47). While we don’t see a decrease in total granular volume in Δ*algP* cells, as might be expected if the protein were critical for phase separation of polyP granules, we do see a change in how polyP is distributed between granules, with fewer, larger granules. We speculate that AlgP may normally function to limit granule fusion (Fig 7). Fusion inhibition might occur by two mechanisms: non-specific or specific interactions of AlgP with DNA when bound to small/nascent polyP granules could restrict their motion such that they can only fuse with local granules tethered within a given chromatin neighborhood (Fig 7A). Such restriction to fusing with local neighbors could lead to even spacing. Another non-exclusive model is that AlgP changes the biophysical properties of granules in a manner that inhibits fusion, such as modulating their surface charge. Future time-lapse imaging experiments of polyP granules are needed to address these models. The fusion behavior of polyP granules in the Δa*lgP* mutant in *P. aeruginosa* differs from the fusion behavior of PHA granules in the Δ*phaF* mutant in another Pseudomonad, *Pseudomonas putida*. In that species and strain, PHA granules bunch together but do not fuse, likely due to the effect of another protein PhaI(36).

**Figure 7.**
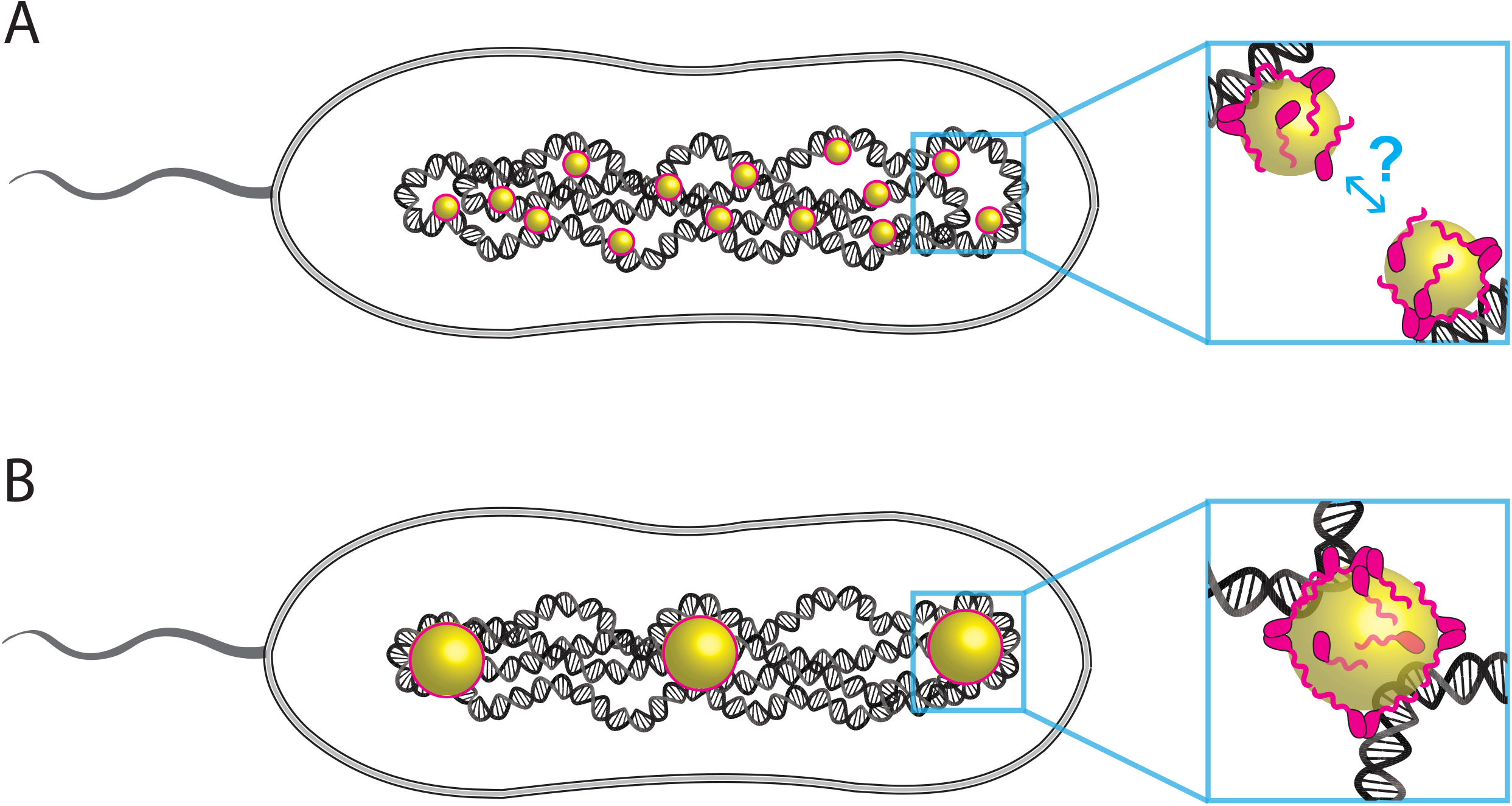
Model for the role of AlgP in granule biogenesis. (A) AlgP tethers nascent polyP granules to the nucleoid, either through specific or non-specific contacts. Interactions between nascent polyP granules and the nucleoid prevent free diffusion of granules, and lead to some degree of local confinement within a chromosomal ‘neighborhood.’ Such confinement allows granules to fuse only with their nearest neighbors on the timescale of granule maturation (hours). (B) Mature granules, with their larger mass and more extensive interactions with the nucleoid, are more limited in their radius of gyration and do not fuse with other granules, leading to even spacing.

### Functional significance of AlgP in starvation fitness and pathogenesis

Having discovered that AlgP modulates granule spacing in cells, the next important question is why cells invest in granule spacing at all. Even spacing of another class of bacterial membraneless compartments, carboxysomes, is important for the fitness of daughter cells and mediated by a Brownian-ratchet mechanism(50, 51). While we don’t yet understand the molecular mechanism fully underpinning AlgP-mediated spacing, we now have the means in *P. aeruginosa* to disrupt polyP granule organization and ask whether even spacing of polyP granules poses either a survival benefit during starvation, or a growth advantage when nutrients become available. The CTD of AlgP has been identified as a hotspot for genetic rearrangements in clinical isolates of *P. aeruginosa* from cystic fibrosis patients(27, 52). These rearrangements alter KPAA repeat number, suggesting that this domain is under selective pressure during chronic infections, however more research is needed to determine whether there is a connection between these rearrangements and infection outcomes. The consequences of these alterations in AlgP CTD domain length for granule biogenesis, starvation fitness, and pathogenesis remain to be explored. Recently an alkyne derivative of the natural product elegaphenone, which has antibiotic effects against *P. aeruginosa*, was shown to bind AlgP, suggesting that AlgP is worthy of further exploration as a possible drug target(53).

## METHODS

### Media and Growth conditions

For nitrogen starvation experiments, strains were grown overnight at 37°C shaking in MOPS-buffered minimal media (MMM): 40mM sodium succinate, 22mM NH_4_Cl, 43mM NaCl, 2.2mM KCl, 1.25mM NaH_2_P0_4_, 1mM MgSO_4_, 0.1mM CaCl_2_, 7.5µM FeCl_2_·4H_2_O, 0.8µM CoCl_2_·6H_2_O, 0.5µM MnCl_2_·4H_2_O, 0.5µM ZnCl2, 0.2µM Na_2_MoO_4_·2H_2_O, 0.1µM NiCl_2_·6H_2_O, 0.1µM H_3_BO_3_, and 0.01µM CuCl_2_·2H_2_O, 50mM MOPS, pH 7.2). Optical density was determined at 500nm, rather than 600nm, to minimize interference of pigments such as phenazines that *P. aeruginosa* produces during starvation. 5mL subcultures at OD_500_ = 0.0125 to 0.025 were grown in glass test tubes at 37°C shaking at 250rpm to OD_500_ = 0.4 to 0.6, then spun down at room temperature for 5 minutes at 5000xG, and resuspended to OD_500_ = 0.4 in nitrogen-limited MMM (Identical to MMM, but with 1mM NH_4_Cl instead of 22mM) in clean glass test tubes and grown at 37°C shaking at 250rpm. Time 0h = cells collected immediately before being spun down and shifted to nitrogen-limited medium.

### Granule isolation and proteomic analysis

Refer to the SI methods for a detailed description of cell culturing (cell growth and harvesting) and proteomic analysis (sample preparation, data acquisition, software used and data availability). Briefly, cells were grown to exponential phase at 37°C in MMM and then shifted to a low-nitrogen MMM for 3h at 37°C. Cell pellets corresponding to 100mL of cell culture were flash frozen in liquid nitrogen. Pellet was resuspended and incubated on ice for 15min in 1mL of lysis buffer, followed by microtip sonication (Qsonica Q700), and additional nucleic acid digestion with Benzonase and DNAse I. Lysate (0.5mL) was loaded onto 7mL pre-chilled Percoll gradient solution in ultracentrifuge tubes and spun in a 50TI rotor at 21,300 RPM for 15 minutes. After removing the gradient solution, a Percoll-encased pellet was resuspended in 1mL of dilution buffer and centrifuged in a tabletop microcentrifuge at 10,000G for 2 min, resuspended, and the spin repeated to remove residual Percoll from the pellet. The pellet was flash frozen in liquid nitrogen for subsequent proteomic analysis. The (g/g) absolute mass fraction of a protein in a given sample was estimated using the spectral counting technique(54). For a protein, its mass fraction abundance was tabulated by dividing the total number its peptide-spectrum matches (PSM) by the total of all 14N PSMs in the sample. Enrichment of a protein in our proteomics screen was defined as the ratio of the abundance of protein in the pellet to the abundance in the lysate. See supplemental methods for more details.

### Plasmids and strains

Strains, plasmids, and primers used in this study are listed in Tables S4a-c respectively. Strain construction was performed as described previously, with details outlined in SI methods(9).

### Fluorescence microscopy

All live cell imaging was acquired with a Nikon Ti2-E inverted microscope. Details of optical setup and acquisition parameters are described in SI methods. Cells were imaged at 26°C on 1% agarose pads containing MMM or MMM lacking ammonium chloride for nitrogen starvation experiments. Agarose pads were imaged inside coverslip bottom dishes with plastic lids (Willco Wells, 0.17 mm/#1.5). For DAPI-stained imaging of polyP granules, live cells were stained for 20 minutes in minimal media with 200µM DAPI from a 10 mM stock in dimethyl sulfoxide. DAPI staining of live cells for the purpose of polyP granule imaging is not 100% efficient, with some cells not taking up enough dye to label granules (to overcome this limitation of DAPI labeling, we used TEM to quantify the fraction of granule-free cells). But DAPI can be used to characterize the number of granules per cell in cells that take up the DAPI dye, and their organization. For Sybr Green staining, cells were stained for 20 minutes at 1X dye in minimal media before imaging.

### Transmission Electron Microscopy (TEM)

Unfixed cells (3µL) were spotted onto glow discharged carbon-coated 200 mesh copper grids with pinpointer grid (Ted Pella 01841-F) for 45 seconds, blotted with Whatman paper, and media salts were washed off of grids by spotting the grids with 3uL of water and rapidly blotting; wash step was repeated 5 times. Samples were analyzed at 120kV with a ThermoFisher Talos L120C transmission electron microscope and images were acquired with a CETA 16M CMOS camera. TEM data was plotted using iqplot: http://dx.doi.org/10.22002/D1.1614.

### Image analysis

Cell segmentation and spot detection for fluorescence microscopy images was performed using MicrobeTracker**(55)**. Spot detection was performed after rolling-ball background subtraction in ImageJ (radius 10 pixels). Demographs of fluorescence intensity were constructed using the method and script developed by Hocking et al., 2012**(56)**. For TEM imaging, granule number and diameter per cell were measured manually using ImageJ as described previously**(9)**.

## Data availability

The mass spectrometry proteomics data have been deposited and will be available upon publication to ProteomeXchange Consortium via the UCSD’s MassIVE repository with the accession codes: MassIVE: MSV000087218 and ProteomeXchange: PXD025444

The raw light and electron microscopy data will be deposited upon publication to zenodo.org

## ACKNLOWLEDGEMENTS

We thank Scott Henderson, Kimberly Vanderpool, and Theresa Fassel of the Core Microscopy Facility at the Scripps Research Institute for their expert assistance in performing the TEM experiments. We thank Megan Bergkessel, Ashok Deniz, Keren Lasker, Michael Manson, Dianne Newman, Daniel Scholl and members of the Racki Lab for helpful feedback and discussions of this work. This work was supported by the Donald E. and Delia B. Baxter Foundation (LRR), and also in part by a grant from the NIH (R35-GM-136412 to JRW). Figures 1A and S1A were created with BioRender.com.

