## Supplementary Information for "The Histone H1-like protein AlgP facilitates even spacing of polyphosphate granules in *Pseudomonas aeruginosa*"

**Supporting Information Appendix**

**SI Figure and table legends**

**SI Methods**

**SI References**

**Table S1.** Summary of proteomics data. Table is included as a separate .csv file

**Table S2a**. Summary of highly abundant and enriched proteins identified in the pellet.

**Table S2b**. Summary of highly enriched proteins and positively charged proteins identified in granules.

**Table S3a.** Fluorescence foci summary.

**Table S3b.** Transmission Electron Microscopy Summary Data.

**Table S3c.** Cell cycle exit.

**Table S4a.** Strains

**Table S4b.** Plasmids

**Table S4c.** Primers

#### SI FIGURE AND TABLE LEGENDS

**FigureS1. Extended proteomic panels of proteome from the three biological experiments shown in Figure 1.** (A) Schematic of polyP granule enrichment protocol. (B) Average absolute mass fraction of proteins in the "pellet" and "lysate", obtained from spectral counting shown in parts per million, as in Fig 1C in the main text, on a linear scale representing the complete and unfiltered proteomic data. (C) Charge of proteins identified in the proteomics analysis plotted against the fold enrichment over the complete charge range (Fig 1E in comparison shows the data for a negative charge cut-off of -75). Figure labels and cut-offs are same as in Figure 1. (D) Absolute mass fraction of proteins in the "pellet" and "lysate", obtained from spectral counting shown in parts per million. Data are shown for a representative experiment. (E) Enrichment of proteins in the pellet plotted against the abundance of proteins in the pellet for a representative experiment. (F) Charge of proteins identified in a representative proteomics experiment plotted against the calculated fold enrichment (negative charge cut-off for plotting: -75). Figure labels and cut-offs are same as in Figure 1.

**Figure S2. AlgP protein sequence.** (A) The boxed region highlights the 154 residue contiguous repeat domain in the C-terminal domain. Highlighted are 25 perfect KPAA repeats (yellow), interspersed with variants, including 7 KPVA (green), 4 KTAAA (cyan), one KPAV, and two alanine spacers, and two KPAA repeats (yellow) that fall outside of this 154aa contiguous region. (B) Full sequence of AlgP as in (A). (C) Alignment of AlgP and PhaF.

**Figure S3. AlgP complementation analysis.** (A) Representative images of DAPI-stained cells. (B) Quantification of the fraction of cells in the population with 1, 2, and >2 DAPI-labeld foci per cell. Each point represents an independent replicate/day of experiment, the bar indicates the mean. Variance analyzed using a one-way ANOVA. Significant differences between strains at the same time point are marked with uppercase letters based on a post hoc Tukey test. Scale bar: 2µm.

**Figure S4. Example of 'satellite' granules in cells imaged by TEM.** (A) Two examples of 3h nitrogen-starved WT cells displaying small satellite granules. (B) Three examples of 0h (exponential phase) WT cells. Top and middle cells display nascent granules, but the bottom cell does not. Scale bar: 0.5µm.

**Figure S5. Effect of polyP and AlgP on growth.** Optical density (500 nm absorbance) as a function of time for *P. aeruginosa* cultures in MOPS minimal media. Cells were subcultured from overnight cultures where cells were preconditioned in MOPS minimal media.

**Table S1. Summary of proteomics data.** The fractional abundance of proteins found in the "pellet" and "lysate" fractions from three independent experiments and their averages are provided in a tabular format.

**Table S2a. Summary of highly abundant and enriched proteins identified in the pellet.** The table summarizes the proteins that were highly abundant in the pellet as well as were highly enriched in the "pellet" fraction. (Abundance cut-off: 5000ppm, Enrichment cut-off:8). Upper right quadrant, Figure 1D

**Table S2b. Summary of highly enriched proteins and positively charged proteins identified in granules.** The table lists the proteins that were both highly abundant in the pellet and highly positively charged. (The enrichment cut-off was 8 fold, the charge cut-off was +5). This table includes the proteins plotted in the upper right quadrant of Fig 1E that have a charge greater than +5.

**Table S3a. Fluorescence foci summary.** Quantification of percentages of 1, 2, and >2-foci cells in DAPI and mApple channels. Values represent average and standard deviation of three independent experiments performed on different days.

**Table S3b. Transmission Electron Microscopy Summary Data.** Average and standard deviation values calculated from the total population of cells as depicted in Figure 4.

**Table S3c. Cell cycle exit.** Quantification of percentage of cells with >1 origin per cell and >0 DNA replication forks per cell. Values represent average and standard deviation of three independent experiments performed on different days, as shown in Figure 6.

**Table S4a. Strains**

**Table S4b. Plasmids**

**Table S4c. Primers**

#### SI METHODS

##### Cell growth and harvesting

*P. aeruginosa* cells were grown as described previously with scale up specific modifications for proteomics noted below(1). WT(PA14) and polyP(LR119) quadruple knockout were streaked from a glycerol stock onto an LB plate and the plate was incubated overnight at 37 °C. A single colony was then inoculated into 25mL complete MOPS minimal media (MMM) and grown to saturation at 37 ºC overnight with shaking at 250 rpm in a 250mL Erlenmeyer flask. Overnight culture (~7mL) was inoculated into 1 L MMM complete media and allowed to grow at 37 ºC to OD500 between 0.4-0.6 in a 2.8L Pyrex Fernbach-style culture flask. The cells were spun down in 1L centrifuge bottles at 5,000 g for 10 min at room temperature (JLA 9.1000 rotor; Avanti J-E centrifuge) and resuspended to an $OD_{500}$ of ~0.4 in 1L low nitrogen MMM media in Pyrex Fernback-style culture flask. At this point, the cultures were incubated back in the 37°C incubator with shaking and allowed to grow for 3h. The cells were then spun in a 1L centrifuge bottle at 5000g for 10min. The supernatant was removed, the pellet resuspended to a volume of 10mL spent medium and aliquoted into Eppendorf tubes (Note:1mL of this cell culture at this step corresponds to 100mL of original cell culture prior to the previous centrifugation step). The Eppendorf tubes were then spun at 5000 g for 5min on a tabletop centrifuge, supernatant discarded and the cell pellets flash frozen in liquid nitrogen. The cell pellets were then stored at -80C till further processing.

##### Granule isolation

Previously generated cell pellets were removed from -80C and incubated on a freshly prepared and chilled lysis buffer. Specifically, the pellet corresponding to each ~200mL cell culture was resuspended in 1mL of lysis buffer and incubated on ice for 15 minutes. The lysis buffer composition was as follows: 200mM NaCl, 35.5mM $Na_2HPO_4$,14.5mM $NaH_2PO_4$, 2mM EDTA, 1X Protease Inhibitor Mix, 0.2mg/mL Lysozyme, 1% Triton X-100. The resuspended solution became extremely viscous upon incubation and was sonicated with microtip sonicator (Qsonica Q700; 70% power, 3 minutes total [10 seconds on, 10 seconds off]) on a CoolRack or ice slurry. After sonication, about ~30µL of this lysate was flash frozen in liquid $N_2$ and stored at -80°C. This fraction was labeled the "lysate" fraction used for proteomic analysis. To the remaining lysate, 1µL of 1M $MgCl_2$, 1µL DNAse I (2000UµL) and 1uL Benzonase (250U/µL) was added per mL of the lysate and the solution was incubated for 30 minutes at room temperature. Next, about ~500uL of the lysate from the previous step was loaded onto 7mL of pre-chilled Percoll gradient solution in ultracentrifuge tubes (Beckman Coulter #355630). Percoll gradient composition was follows: 90% Percoll (Cytiva #17089101), 200 mM NaCl, 35.5mM $Na_2HPO_4$, 14.5mM $NaH_2PO_4$ and 3mM $MgCl_2$. Ultracentrifugation was performed on a pre-chilled rotor (50TI) at 21,300 rpm for 15 minutes in an ultracentrifuge (Beckman Coulter Optima L-80 XF) maintained at 4°C. Upon ultracentrifugation a percoll-encased polyP pellet was clearly visible in the bottom of the tube for wildtype cells and absent in the case of quadruple polyP mutant. The supernatant was removed from the ultracentrifuge tube using a serological pipette leaving behind a Percoll-encased pellet. The Percoll-encased pellet was resuspended in 1mL of dilution buffer (200 mM NaCl, 35.5mM $Na_2HPO_4$, 14.5mM $NaH_2PO_4$, 3mM $MgCl_2$). The resuspended sample was then centrifuged in a table-top microcentrifuge at 10,000 g for 2min and the supernatant discarded. The pellet was then resuspended in 200uL of dilution buffer and washing was repeated once more to remove the Percoll. The samples at this step were labeled as "pellet" samples, flash frozen in liquid $N_2$ and stored at -80°C.

**Proteomics Sample Preparation**
Percoll pellet biomass was resuspended in 100 µL HPLC-grade water and protein was precipitated by methanol–chloroform extraction(2). Insoluble material was pelleted by centrifugation at 17,000 *g* and re-solubilized in 40 µL of freshly-made 8M urea buffered by 100 mM Tris-HCl, pH 7.5. Disulfides were reduced with 10 mM DTT (30 min at 37°C), alkylated with 40 mM iodoacetamide (30 min at 37C). Reaction was quenched with 20 mM DTT (30 min at 37°C).

For compatibility with tryptic digestion, reaction volume was diluted with 50 mM Tris-HCl, 10 mM $CaCl_2$ to 200 µL (final concentration of 1.6M Urea). Cellular protein was digested for 16 hours at 37C with 2 µg porcine pancreatic trypsin (Thermo), and followed by an booster dose of 1 ug trypsin for 4 hours. Peptides were de-salted using PepClean columns (Thermo), dried using a SpeedVac, and re-suspended in 20 µL of LC-MS solvent A (Honeywell, 0.1% formic acid in water) prior to analysis.

**Proteomics Data Acquisition**
Tryptic peptides (2-3 µL, approximately 1-2 µg based on absorbance) were injected onto an Eksigent EKSPERT NanoLC 425 chromatography system operating in trap-elute mode. Peptides were eluted from a SCIEX ChromXP analytical C18 reverse-phase nanoflow column (3 µm, 120Å, 150 x 0.3 mm) over the course of a 120-minute linear ramp gradient (5-35% of 0.1% formic acid in acetonitrile) at 300 nL/min. A SCIEX TripleTOF 5600 operating in DDA mode was used for data acquisition in positive polarity mode. An MS1 survey scan (250 ms, 400-1250 Th, high-resolution) was followed by 20 product ion scans (150 ms, 100-1500 Th, high-sensitivity). Ions with a charge of +2 to +4 exceeding 100 counts-per-second were subjected to fragmentation. Collision energy was set to 'rolling'. Former targets ions were excluded for 15 sec after one occurrence.

**Proteomics Data Analysis**
Vendor (.WIFF format) data files were converted with the SCIEX MS Data Converter (Beta 1.3) to mzXML format in centroid mode. The trans-Proteomic Pipeline software suite was used to search the data with X!TANDEM against a UniProt *Pseudomonas aeruginosa (strain UCBPP-PA14)* database (proteome UP000000653, retrieved October 2019) supplemented with common contaminants, enzymes and reversed peptide decoy sequences(3, 4). The peptide-spectrum match tolerances were: 50 ppm and 100 ppm for the precursor and product ions. PeptideProphet and iProphet were used to combine the peptide–spectrum matches across multiple samples and SpectraST was used to generate spectral libraries and collate search results(5).

*Absolute quantification.* To estimate the (g/g) absolute mass fraction of a protein in a given sample, we used the spectral counting technique(6). For a protein, its mass fraction abundance was tabulated by dividing the total number its peptide-spectrum matches (PSM) by the total of all 14N PSMs in the sample. The abundances of proteins from three independent experiments (and the averages, refer to the section below) are reported in Table S1.

**Software used for analyses.** Further proteomic data processing and analysis was performed in Python (CPython 3.7.7, IPython 7.21.0) with NumPy version 1.19.2 and Pandas version 1.2.3 using Jupyter notebook (Jupyerlab version 3.0.11). Data was plotted with Bokeh version 2.3.0 and the figures were assembled in Adobe Illustrator. Enrichment of a protein in our proteomics screen was defined as the ratio of the abundance of protein in the pellet to the abundance in the lysate. This definition leads to an "infinite" enrichment when abundance of a protein in the lysate is zero and to computationally handle these infinite values in Pandas DataFrame a value of 512 ($2^9$ i.e., $log_2$ fold change=9) was assigned. The averaging of proteins from three separate experiments was performed using the built-in mean function of the Pandas Dataframe with the 'skipna' parameter set to 'True' to exclude the NA/null values when computing the result. An outcome of this selection is that the proteins with infinite enrichment in one or more experiments show up as a band or clusters, an artifact we termed "banding." In our current analysis we have not pursued the banding pattern, but in future it would be curious to probe if some of the proteins exhibiting the banding pattern could be part of the interactome of the polyphosphate granule that transiently and/or weakly associated. The charges of the protein (at pH 7) on PA14 proteome were obtained from UniPort (UP000014183; date accessed February 18th, 2020) and integrated it in Jupyter notebook analysis pipeline by a Panda DataFrame merge operation on the protein locus ID. An abundance value of >5000 ppm and enrichment value of 8 ($2^3$ i.e., $log_2$ fold change = 3) were, respectively, used as cut-offs for "high" abundance and enrichment in our analysis discussed in Figure 1 (also see Table S2A,B).

**Fluorescence microscopy**
All live cell imaging was acquired with a Nikon Ti2-E inverted microscope with perfect focus and the following other hardware: Objective: Plan apochromat phase contrast 100X oil immersion objective, N.A. 1.45, Illumination Source: For brightfield, a white LED, for fluorescence, the Spectra X Light Engine with a 470nm LED (Lumencor). Camera: Prime 95B sCMOS with 11 µm x 11 µm pixel area (Photometrics). Image acquisition was controlled using Nikon Elements. The following parameters were used: For phase contrast: 75% light intensity, 100ms exposure time. For mNeon, GFP, and SybrGreen imaging: 100% light intensity from the 470nm LED, 100ms exposure time and a GFP filter cube (466/40nm excitation filter, 525/50nm emission filter, 495nm dichroic mirror, Semrock). For mCherry and mApple: 100% light intensity from the 555nm LED, 100ms exposure time and Texas Red filter cube (562/40nm excitation filter, 641/75nm emission filter, 593nm dichroic mirror, Semrock). For DAPI imaging of polyP: 100% light intensity from the 395nm LED, 100ms exposure time and a custom filter cube (415/20nm excitation filter, 555/10nm emission filter, 425nm long pass dichroic mirror, Semrock).

**Data Availability.**
The mass spectrometry proteomics data have been deposited and available to ProteomeXchange Consortium via the UCSD's MassIVE repository with the accession codes: MassIVE: MSV000087218 and ProteomeXchange: PXD025444.

**Strain construction**
Strains, plasmids, and primers used in this study are listed in Tables S7-9 respectively.

**Strains**
All unmarked deletion strains, and strains in which endogenous proteins are replaced by fluorescent chimeras, were generated by triparental conjugation with *P. aeruginosa* UCBPP-PA14, and then merodiploids were selected as described previously on VBMM medium (3 g/L trisodium citrate, 2 g/L citric acid, 10g/L $K_2HO_4$, 3.5 g/L $NaNH_4PO_4$, 1mM $MgSO_4$, 100uM $CaCl_2$, pH 7) containing 100 ug/mL gentamicin(7). Counterselection for homologous recombination events removing the endogenous copy of the gene in question was then performed on LB plates without NaCl and containing 300mM sucrose, followed by PCR verification. All strains with insertions at the *att*Tn7 site were generated by tetraparental conjugation with *P. aeruginosa* UCBPP-PA14, and then exconjugants were selected on

VBMM medium, and verified by PCR(7).

**Plasmids**
All plasmids were generated using either Gibson cloning or the yeast gap repair method of homologous recombination by *Saccharomyces cerevisiae*(8–10). Inserts were generated by PCR, or from synthetic gBlock gene fragments (Integrated DNA Technologies). Plasmids pLREX79, pLREX120, pLREX120, pLREx121, pLREX124, and pLREX125 are derivatives of suicide vector pMQ30 (2), generated by amplifying ~1 kb of sequence upstream and downstream of the target gene from *P. aeruginosa* genomic DNA. Plasmid pLREX132 is a derivative of the pUC18T-mini-Tn7T-Gm suicide vector.

**pLREX79** *[ppk2A::ppk2A-20aa-mNeonGreen]* was created by yeast homologous recombination between digested plasmid pLREX9 and gBlock6. Plasmid pLREX9 [*ppk2A::ppk2A-mCherry*] was cut with NotI and XmaI to remove mCherry. Plasmid confirmed by Sanger sequencing.

**pLREX120** [*ΔalgP*] was created by a Gibson assembly consisting of pMQ30 cut with HindIII and KpnI and 2 fragments: (1) 815bp PCR product of template *P. aeruginosa* PA14 genomic DNA, primers LRPR894F and LRPR912R, (2) 540bp PCR product of template *P. aeruginosa* PA14 genomic DNA, primers LRPR909F and LRPR899R. Plasmid confirmed by Sanger sequencing.

**pLREX121** [*algPΔCTD*] was created by Gibson assembly consisting of pMQ30 cut with HindIII and KpnI and 2 fragments: (1) 1286bp PCR product of template *P. aeruginosa* PA14 genomic DNA, primers LRPR894F and LRPR933R, (2) 543bp PCR product of template *P. aeruginosa* PA14 genomic DNA, primers LRPR932F and LRPR899R. Plasmid confirmed by Sanger sequencing.

**pLREX124** [*algP::mApple-algP*] was created by Gibson assembly consisting of pMQ30 cut with HindIII and KpnI and 3 fragments: (1) 818bp PCR product of template *P. aeruginosa* PA14 genomic DNA, primers LRPR894F and LRPR904R, (2) 772bp PCR product of template gBlock4, primers LRPR905F and LRPR906R, and (3) 1594bp PCR product of template *P. aeruginosa* PA14 genomic DNA, primers LRPR907F and LRPR899R. Plasmid confirmed by Sanger sequencing.

**pLREX125** [*algP::algP-mApple*] was created by Gibson assembly consisting of pMQ30 cut with HindIII and KpnI and 3 fragments: (1) 1886bp PCR product of template *P. aeruginosa* PA14 genomic DNA, primers LRPR894F and LRPR901R, (2) 771bp PCR product of template gBlock4, primers LRPR900F and LRPR903R, and (3) 535bp PCR product of template *P. aeruginosa* PA14 genomic DNA, primers LRPR898F and LRPR899R. Plasmid confirmed by Sanger sequencing.

**pLREX132** [ $P_{algP}$:*algP*] was created by Gibson assembly consisting of pUC18T-mini-Tn7T-Gm cut with HindIII and KpnI and 1 fragment: a 1181bp PCR product of template *P. aeruginosa* PA14 genomic DNA and primers LRPR956F and LRPR955R. This fragment contains the *algP* coding sequence and the 125 bp intergenic region upstream of algP containing the putative previously identified promoter sequence 'CGAACCCGTTGGCGAGAGGGGGTTTGCGGGTCTAGTATGGGCGCAACCAC' from *P. aeruginosa* PA14 genomic DNA(11, 12).

**Sequences of gBlocks, fluorescent proteins, and linkers**

**gBlock4:**
(***bold/italicized sequence is mApple,*** GenBank: DQ336160.2, codon optimized for *Pseudomonas aeruginosa*)
CGAGGAGGACGAGAAGGTCTACGCCGAGGCGGCCGCCGCGCCGGGCCACGCGAACCT
GGATATCCCGGCCCTCGAGGGGTCCGGTCAGGGACCGGGATCCGGCCAAGGGTCCGG
C***ATGGTGTCGAAGGGCGAGGAAAACAATATGGCCATCATCAAAGAGTTCATGCGGTTC***
***AAGGTCCATATGGAGGGGTCGGTCAATGGGCACGAGTTCGAGATCGAAGGCGAAGGC***
***GAGGGGCGGCCGTATGAGGCGTTCCAGACCGCGAAGCTGAAGGTCACGAAGGGGGGG***
***CCGCTCCCCTTCGCGTGGGACATCCTCTCCCCCCAATTCATGTATGGCTCCAAAGTCTA***
***CATCAAGCATCCGGCCGATATCCCCGATTATTTCAAGCTGAGCTTCCCCGAGGGCTTCC***
***GCTGGGAACGGGTCATGAATTTCGAAGATGGCGGGATCATCCACGTGAACCAAGATAG***
***CAGCCTCCAAGATGGCGTGTTCATCTATAAGGTCAAACTGCGCGGGACGAATTTCCCCT***
***CCGATGGGCCCGTGATGCAGAAAAAAACGATGGGCTGGGAAGCCAGCGAGGAACGCA***
***TGTATCCCGAAGACGGCGCCCTGAAGTCGGAGATCAAAAAACGGCTCAAGCTGAAAGA***
***TGGGGGCCACTATGCGGCGGAGGTGAAAACCACGTACAAGGCGAAAAAGCCCGTGCA***
***ACTCCCCGGCGCGTACATCGTGGACATCAAACTCGATATCGTGAGCCACAATGAGGAC***
***TATACCATCGTCGAGCAGTATGAACGCGCCGAAGGCCGGCACTCGACCGGGGGGATG***
***GACGAACTCTATAAATAA***GGCGGGCGGTCGCGCCAACGAAAACGCCCGGGGCGCTTTC
GCGCTCCGGGCGTCCC

**gBlock6:**
(underlined sequence is 20aa Linker 1, ***bold/italicized sequence is mNeonGreen)***
CGAGGAGGACGAGAAGGTCTACGCCGAGGCGGCCGCCGCGCCGGGCCACGCGAACCT
GGATATCCCGGCCGGATCCGGGCAGGGACCGTCTGGCCAGGGATCGGGGCCAGGATCA
GGTCAAGGCTCCGGT***ATGGTTTCGAAAGGAGAGGAGGATAATATGGCTAGCCTCCCAG***
***CGACCCACGAACTGCATATTTTTGGCAGCATTAATGGCGTTGACTTTGATATGGTGGGG***
***CAGGGAACAGGGAACCCTAACGATGGCTATGAGGAGCTCAATCTCAAGAGTACAAAAG***
***GAGATTTGCAATTTTCACCTTGGATCCTGGTTCCGCATATTGGCTACGGCTTTCATCAAT***
***ACTTGCCTTATCCGGACGGCATGTCCCCGTTCCAAGCTGCGATGGTGGATGGTTCTGGG***
***TACCAGGTGCACCGTACTATGCAGTTTGAGGACGGTGCCTCACTGACGGTCAACTATAG***
***ATATACTTATGAAGGCTCACACATTAAGGGTGAGGCCCAAGTTAAAGGAACAGGGTTTC***
***CTGCGGATGGACCGGTAATGACAAACAGTTTAACCGCTGCGGACTGGTGTCGCTCGAA***
***AAAAACATACCCAAACGATAAAACGATCATCTCGACCTTCAAATGGAGCTATACTACGG***
***GCAACGGCAAACGCTATCGTTCCACAGCACGCACGACTTATACGTTTGCTAAACCGATG***
***GCCGCAAACTACCTCAAAAATCAACCTATGTACGTGTTCAGAAAAACCGAGTTAAAACA***
***TTCAAAAACGGAACTTAATTTTAAAGAGTGGCAAAAGGCGTTTACAGACGTGATGGGTA***
***TGGATGAACTCTATAAGTGA***GGCGGGCGGTCGCGCCAACGAAAACGCCCGGGGCGCTT
TCGCGCTCCGGGCGTCCC

**mNeonGreen:**
ATGGTTTCGAAAGGAGAGGAGGATAATATGGCTAGCCTCCCAGCGACCCACGAACTGCA
TATTTTTGGCAGCATTAATGGCGTTGACTTTGATATGGTGGGGCAGGGAACAGGGAACCC
TAACGATGGCTATGAGGAGCTCAATCTCAAGAGTACAAAAGGAGATTTGCAATTTTCACCT
TGGATCCTGGTTCCGCATATTGGCTACGGCTTTCATCAATACTTGCCTTATCCGGACGGC
ATGTCCCCGTTCCAAGCTGCGATGGTGGATGGTTCTGGGTACCAGGTGCACCGTACTAT
GCAGTTTGAGGACGGTGCCTCACTGACGGTCAACTATAGATATACTTATGAAGGCTCACA
CATTAAGGGTGAGGCCCAAGTTAAAGGAACAGGGTTTCCTGCGGATGGACCGGTAATGA
CAAACAGTTTAACCGCTGCGGACTGGTGTCGCTCGAAAAAAACATACCCAAACGATAAAA
CGATCATCTCGACCTTCAAATGGAGCTATACTACGGGCAACGGCAAACGCTATCGTTCCA

CAGCACGCACGACTTATACGTTTGCTAAACCGATGGCCGCAAACTACCTCAAAAATCAAC
CTATGTACGTGTTCAGAAAAACCGAGTTAAAACATTCAAAAACGGAACTTAATTTTAAAGA
GTGGCAAAAGGCGTTTACAGACGTGATGGGTATGGATGAACTCTATAAGTGA

**mApple:**
ATGGTGTCGAAGGGCGAGGAAAACAATATGGCCATCATCAAAGAGTTCATGCGGTTCAAG
GTCCATATGGAGGGGTCGGTCAATGGGCACGAGTTCGAGATCGAAGGCGAAGGCGAGG
GGCGGCCGTATGAGGCGTTCCAGACCGCGAAGCTGAAGGTCACGAAGGGGGGGCCGCT
CCCCTTCGCGTGGGACATCCTCTCCCCCCAATTCATGTATGGCTCCAAAGTCTACATCAA
GCATCCGGCCGATATCCCCGATTATTTCAAGCTGAGCTTCCCCGAGGGCTTCCGCTGGG
AACGGGTCATGAATTTCGAAGATGGCGGGATCATCCACGTGAACCAAGATAGCAGCCTC
CAAGATGGCGTGTTCATCTATAAGGTCAAACTGCGCGGGACGAATTTCCCCTCCGATGGG
CCCGTGATGCAGAAAAAAACGATGGGCTGGGAAGCCAGCGAGGAACGCATGTATCCCGA
AGACGGCGCCCTGAAGTCGGAGATCAAAAAACGGCTCAAGCTGAAAGATGGGGGCCACT
ATGCGGCGGAGGTGAAAACCACGTACAAGGCGAAAAAGCCCGTGCAACTCCCCGGCGC
GTACATCGTGGACATCAAACTCGATATCGTGAGCCACAATGAGGACTATACCATCGTCGA
GCAGTATGAACGCGCCGAAGGCCGGCACTCGACCGGGGGGATGGACGAACTCTATAAAT
AA

**20aa linker 1**
ggatccgggcagggaccgtctggccagggatcggggccaggatcaggtcaaggctccggt

**Figure S1**

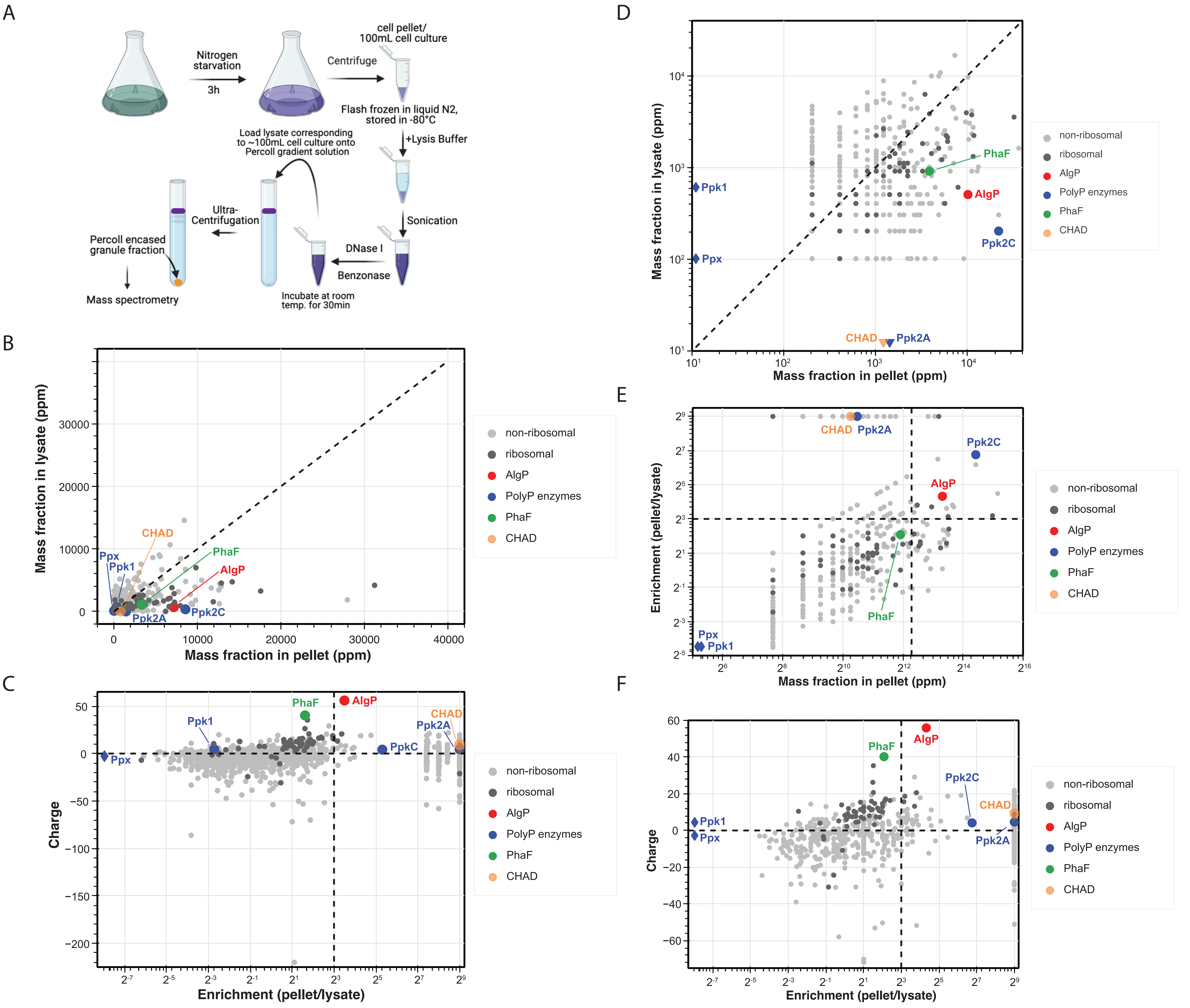
A
Nitrogen starvation
3h
Centrifuge
cell pellet/ 100mL cell culture
Flash frozen in liquid N2, stored in -80°C
+Lysis Buffer
Sonication
DNase I
Benzonase
Incubate at room temp. for 30min
Load lysate corresponding to ~100mL cell culture onto Percoll gradient solution
Ultra-Centrifugation
Percoll encased granule fraction
Mass spectrometry
B
Mass fraction in lysate (ppm)
Mass fraction in pellet (ppm)
CHAD
PhaF
AlgP
Ppx
Ppk1
Ppk2A
Ppk2C
C
Charge
Enrichment (pellet/lysate)
PhaF
AlgP
Ppk1
CHAD
Ppk2A
PpkC
Ppx
D
Mass fraction in lysate (ppm)
Mass fraction in pellet (ppm)
Ppk1
Ppx
PhaF
AlgP
Ppk2C
CHAD
Ppk2A
E
Enrichment (pellet/lysate)
Mass fraction in pellet (ppm)
CHAD
Ppk2A
Ppk2C
AlgP
PhaF
Ppx
Ppk1
F
Charge
Enrichment (pellet/lysate)
AlgP
PhaF
Ppk2C
CHAD
Ppk1
Ppx
Ppk2A
non-ribosomal
ribosomal
AlgP
PolyP enzymes
PhaF
CHAD

### Figure S2

A

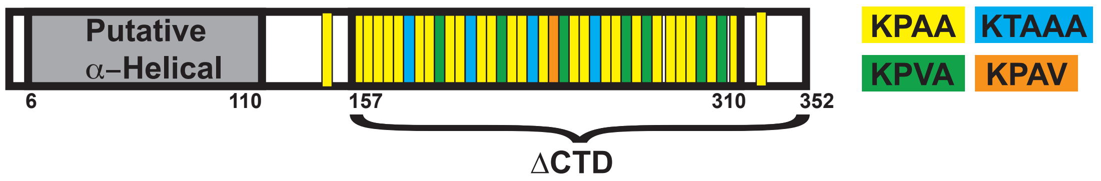


B

MSANKKPVTTPLHLLQQLSHSLVEHLEGACKQALV
DSEKLLAKLEKQRGKAQEKLHKARTKLQDAAKAG
KTKAQAKARETISDLEEALDTLKARQADTRTYIVGL
KRDVQESLKLAQGVGKVKEAAGKALESRKAKPAT
KPAAKAAAKPAMKTVAAKPAAKPAAKPAAKPAAK
PAAKTAAAKPAAKPAAKPVAKPAAKPAAKTAAAK
PAAKPAAKPVAKPAAKPAAKTAAAKPAAKPAVKP
VAKPAAKPAAKTAAAKPAAKPAAKPVAKPAAKPV
AKPAAAKPAAKPAAKPAAKPVAKPAAKPVAAKPA
AAKPATAPAAKPAATPSAPAAASSAASATPAAGSN
GAAPTSAS

C

```
AlgP    1 MSANKKPVTT--PLHLLQQLSHSLVEHLEGACKQALVDSEKLLAKLEKQRGKAQEKLHKA
PhaF    1 MAGKKKSEKESSWIGEIEKYSRQIWLAGLGAYSKVSKDGSKLFETLVKDGEKAEKEA-KS

AlgP   59 RTKLQ-DAAKAGKTKAQAKAR-------ETISDLEEALDTLKARQADTRTYIVGLKRDVQ
PhaF   60 DVDAQVGAAKASARSAKSKVDEVRDRALGKWSELEEAFDKR-LNSAISRLGVPS------

AlgP  111 ESLKLAQGVGKVKEAAGKALE-SRKAKPATKPAAKAAAKPAMKTVAAKPAAKPAAKPAAK
PhaF  113 --------RNEVKELHGKVDTLTRQI-------------EKLTGVSVKPAAKAAAKPAAK

AlgP  170 PAAKPAAKTAAAKPAAKPAAKPVAKPAAKPAAKTAAAKPAAKPAAKPVAKPAAKPAAKTA
PhaF  152 PAAKPAAKTAAAKPAAKPAAKAAAKPAAKPAAKKPAAKTA---AAKPAAKPAAKPAAK-A

AlgP  230 AAKPAAKPAVKPVAKPAAKPAAKTAAAKPAAKPAAKPVAKPAAKPVAKPAAAKPAAKPAA
PhaF  208 AAKPAAKPAAKAAAKPAAKPAAAKPAAKPAAKPAAATAAKPAAKPAAKPAAKKPAAK---

AlgP  290 KPAAKPVAKPAAKPVAAKPAAAKPATAPAAKPAATPSAPAAASSAASATPAAGSNGAAPT
PhaF  265 --------KPAAKPAAAKPAAPAASSSAPAAPAATPAASAPAANA-PA---------TPS

AlgP  350 SAS
PhaF  307 SQG
```

### Figure S3

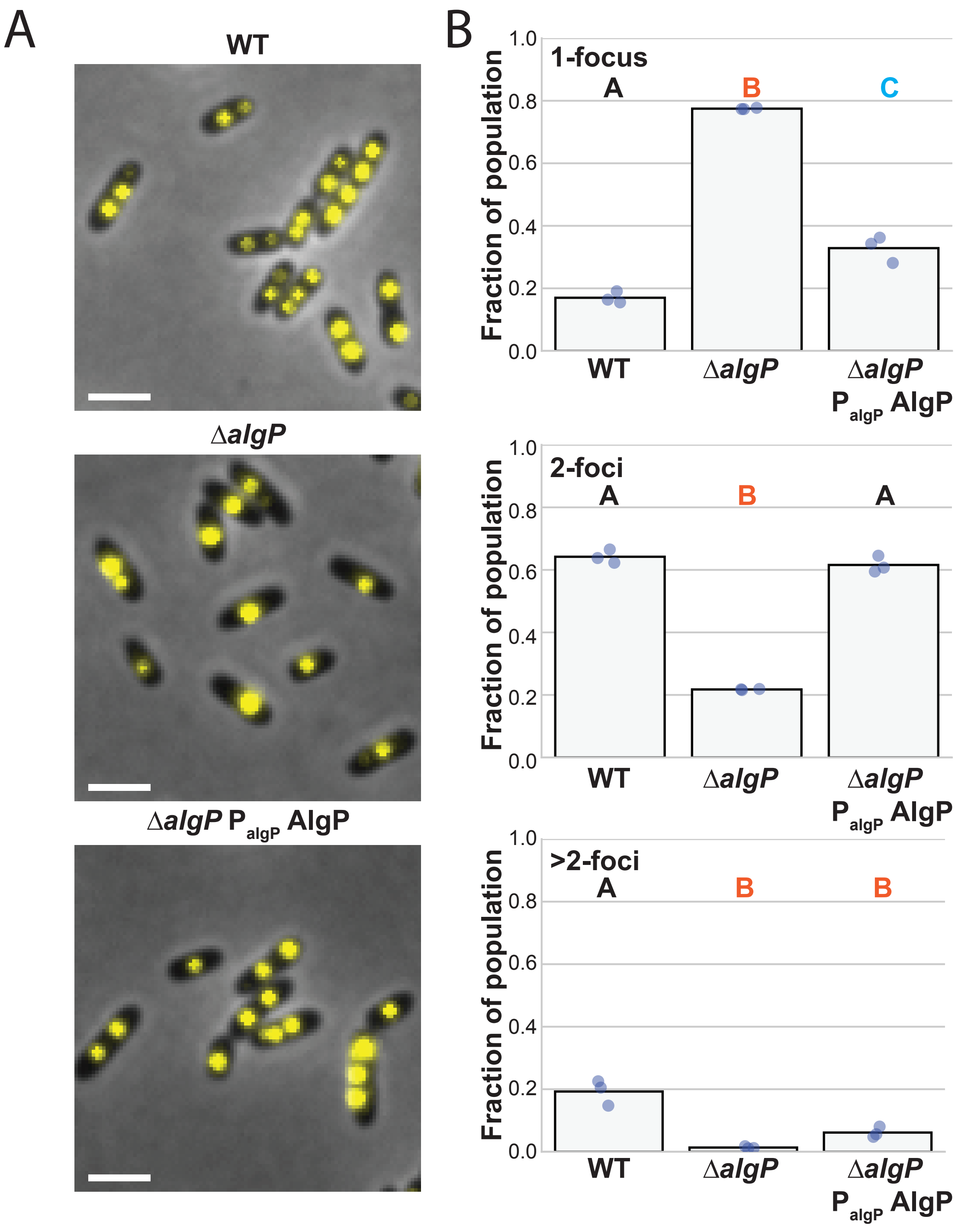
A
WT
ΔalgP
ΔalgP P algP AlgP
B
1-focus
2-foci
>2-foci
A
B
C
Fraction of population
0.0
0.2
0.4
0.6
0.8
1.0
WT
ΔalgP
ΔalgP P algP AlgP

**Figure S4**

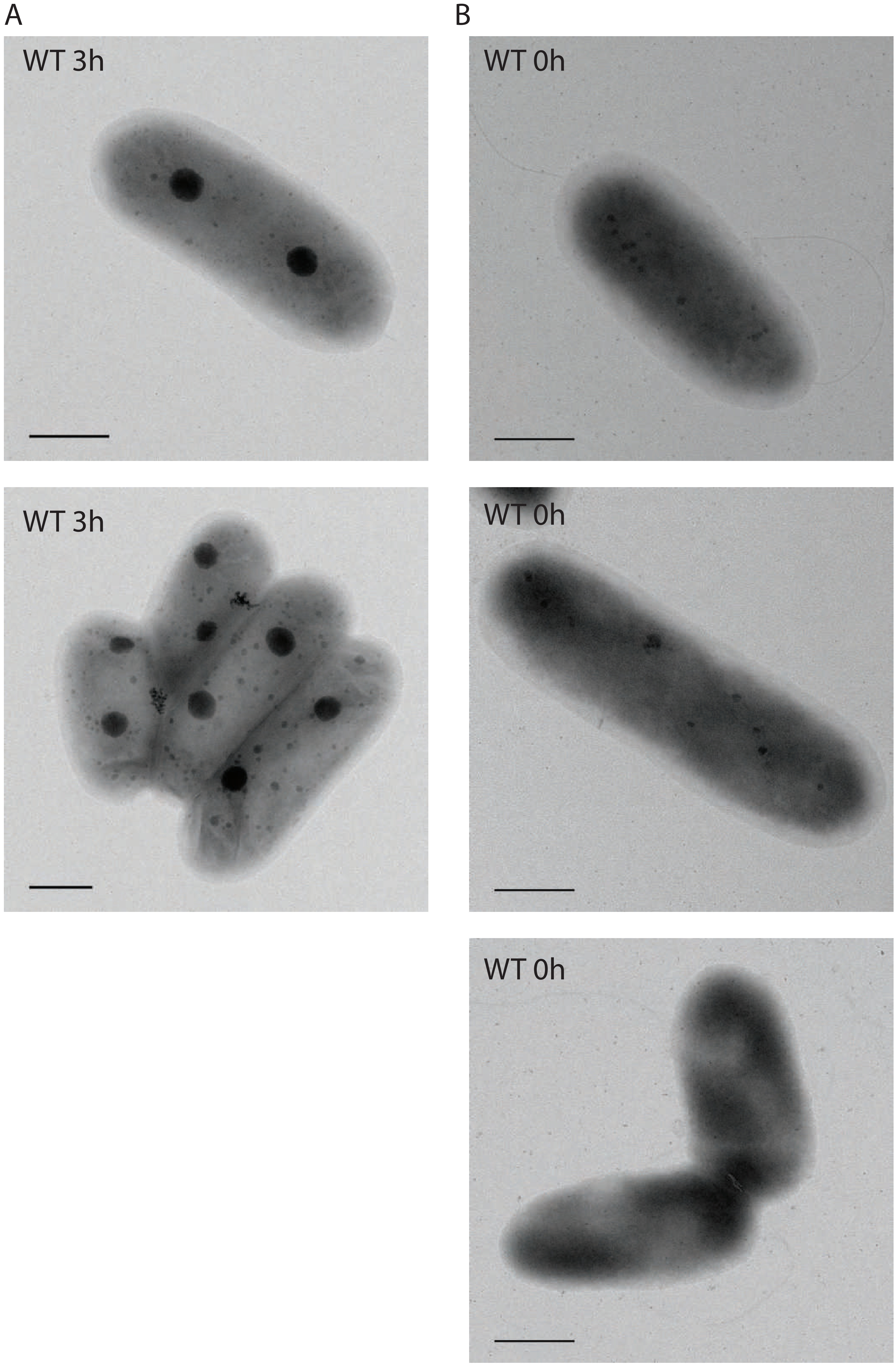

**Figure S5**

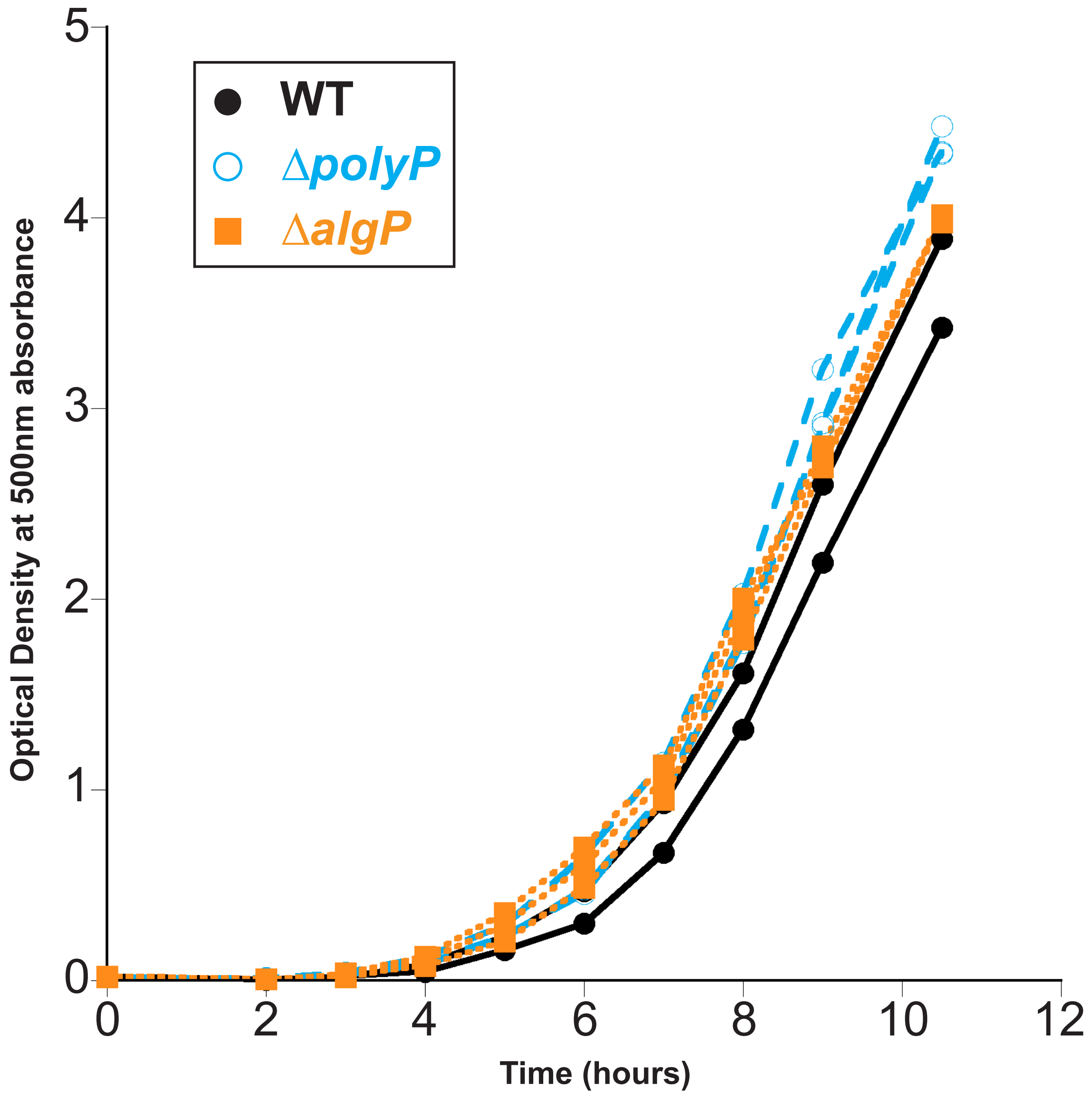

WT
ΔpolyP
ΔalgP
Optical Density at 500nm absorbance
Time (hours)
0
1
2
3
4
5
0
2
4
6
8
10
12

#### Table S2a: Summary of highly abundant and enriched proteins identified in the pellet

| Locus tag | Gene | Description | Average Fraction abundance (pellet) | Average Enrichment (Pellet/Lysate) | Estimated Charge (pH 7) |
|---|---|---|---:|---:|---:|
| **PA14_07560** | rpsU | 30S ribosomal protein S21 | 6598.21683 | 8.958031449 | 13.19 |
| **PA14_08790** | rpsL | 30S ribosomal protein S12 | 7938.981943 | 10.31125836 | 20.61 |
| **PA14_19410** | ppk2C | Polyphosphate kinase | 8551.646369 | 39.5507778 | 4.14 |
| **PA14_29590** | Unknown | Putative transcriptional regulator | 9409.912338 | 8.903982375 | 5.47 |
| **PA14_49740** | Unknown | Uncharacterized protein | 6244.541078 | 222.9901809 | 6.83 |
| **PA14_56070** | mvaT | Transcriptional regulator MvaT, P16 subunit | 27969.53852 | 15.482892 | 6.47 |
| **PA14_65200** | rnr | Ribonuclease R (RNase R) (EC 3.1.13.1) | 12620.7254 | 27.19592607 | 19.14 |
| **PA14_68660** | rimK | Probable alpha-L-glutamate ligase (EC 6.3.2.-) | 6165.75888 | 512 | 8.83 |
| **PA14_69370** | algP | Alginate regulatory protein AlgP | 7151.683224 | 11.27486611 | 55.91 |

**Table S2b: Summary of highly abundant and enriched proteins identified in the pellet**

| Locus tag | Gene | Description | Average Enrichment (Pellet/Lysate) | Estimated Charge (pH 7) |
|---|---|---|---|---|
| **PA14_69370** | algP | Alginate regulatory protein AlgP | 11.2748661 | 55.91 |
| **PA14_12760** | Unknown | Probable ATP-dependent RNA helicase | 350.866132 | 21.87 |
| **PA14_19290** | srmB | Putative ATP-dependent RNA helicase | 186.46567 | 21.09 |
| **PA14_08790** | rpsL | 30S ribosomal protein S12 | 10.3112584 | 20.61 |
| **PA14_65080** | ygiR | UPF0313 protein PA14_65080 | 512 | 19.74 |
| **PA14_66210** | waaX | Putative lipopolysaccharide core biosynthesis protein | 256.5296 | 19.35 |
| **PA14_65200** | rnr | Ribonuclease R (RNase R) (EC 3.1.13.1) | 27.1959261 | 19.14 |
| **PA14_16860** | plsB | Glycerol-3-phosphate acyltransferase (GPAT) (EC 2.3.1.15) | 172.078933 | 19 |
| **PA14_15350** | Unknown | Putative integrase | 19.3861906 | 18.67 |
| **PA14_25000** | slt | Putative soluble lytic transglycosylase | 256.5296 | 18.29 |
| **PA14_66720** | priA | Primosomal protein N' (EC 3.6.4.-) (ATP-dependent helicase PriA) | 512 | 16.62 |
| **PA14_64490** | Unknown | Rho_N domain-containing protein | 512 | 16.47 |

| | | | | |
|---|---|---|---|---|
| **PA14_37820** | Unknown | DUF2235 domain-containing protein | 512 | 14.51 |
| **PA14_66190** | Unknown | Uncharacterized protein | 175.256533 | 14.3 |
| **PA14_63060** | smpB | SsrA-binding protein (Small protein B) | 512 | 13.85 |
| **PA14_53590** | Unknown | Uncharacterized protein | 171.725866 | 13.33 |
| **PA14_07560** | rpsU | 30S ribosomal protein S21 | 8.95803145 | 13.19 |
| **PA14_69710** | xerC | Tyrosine recombinase XerC | 512 | 13.1 |
| **PA14_05970** | Unknown | Uncharacterized protein | 171.019733 | 12.86 |
| **PA14_73420** | rnpA | Ribonuclease P protein component (RNase P protein) (RNaseP protein) (EC 3.1.26.5) (Protein C5) | 512 | 12.19 |
| **PA14_11080** | cupB3 | Usher CupB3 | 512 | 11.55 |
| **PA14_25450** | lolE | Putative lipoprotein releasing system, permease protein | 512 | 11.38 |
| **PA14_57340** | murG | UDP-N-acetylglucosamine--N-acetylmuramyl-(pentapeptide) pyrophosphoryl-undecaprenol N-acetylglucosamine transferase (EC 2.4.1.227) (Undecaprenyl-PP-MurNAc-pentapeptide-UDPGlcNAc GlcNAc transferase) | 256 | 11.25 |
| **PA14_23410** | orfJ | Putative glycosyl transferase | 180.542645 | 11.22 |

| | | | | |
|---|---|---|---|---|
| **PA14_24665** | Unknown | THUMP domain-containing protein | 256 | 11.01 |
| **PA14_00060** | Unknown | Putative acyltransferase | 512 | 10.86 |
| **PA14_08440** | Unknown | Putative short chain alcohol dehydrogenase | 512 | 10.63 |
| **PA14_24620** | Unknown | Uncharacterized protein | 512 | 10.27 |
| **PA14_10160** | fepD | Ferric enterobactin transport protein FepD | 512 | 9.89 |
| **PA14_22270** | Unknown | Possible recombinase | 341.333333 | 9.48 |
| **PA14_56920** | inaA | InaA protein | 512 | 9.34 |
| **PA14_26940** | Unknown | CHAD domain-containing protein | 512 | 9.32 |
| **PA14_40290** | lasA | Protease LasA (EC 3.4.24.-) (Staphylolytic protease) | 512 | 9.26 |
| **PA14_14160** | Unknown | Putative acetyltransferase | 512 | 9.03 |
| **PA14_48115** | aprD | Alkaline protease secretion protein AprD | 512 | 8.88 |
| **PA14_12770** | Unknown | Uncharacterized protein | 512 | 8.86 |
| **PA14_68660** | rimK | Probable alpha-L-glutamate ligase (EC 6.3.2.-) | 512 | 8.83 |
| **PA14_14040** | rhlB | ATP-dependent RNA helicase RhlB (EC 3.6.4.13) | 10.006317 | 8.59 |
| **PA14_31560** | Unknown | Putative transcriptional regulator, LysR family | 172.785066 | 8.58 |

| | | | | |
|---|---|---|---|---|
| **PA14_32460** | Unknown | Putative transcriptional regulator | 512 | 8.52 |
| **PA14_30710** | Unknown | RNA chaperone ProQ | 256 | 8.42 |
| **PA14_46020** | Unknown | DTW domain-containing protein | 512 | 8.29 |
| **PA14_66150** | Unknown | Uncharacterized protein | 512 | 8.18 |
| **PA14_30660** | uvrC | UvrABC system protein C (Protein UvrC) (Excinuclease ABC subunit C) | 512 | 8.16 |
| **PA14_66920** | ubiB | Probable protein kinase UbiB (EC 2.7.-.-) (Ubiquinone biosynthesis protein UbiB) | 11.7863626 | 8.11 |
| **PA14_64280** | Unknown | Putative branched-chain amino acid ABC transporter, permease protein | 256.5296 | 7.57 |
| **PA14_04810** | Unknown | Aldehyde dehydrogenase | 341.333333 | 7.35 |
| **PA14_64000** | Unknown | Putative translation initiation factor SUI1 | 512 | 7.31 |
| **PA14_06540** | bioC | Malonyl-[acyl-carrier protein] O-methyltransferase (Malonyl-ACP O-methyltransferase) (EC 2.1.1.197) (Biotin synthesis protein BioC) | 512 | 7.28 |
| **PA14_39410** | Unknown | Putative acetyltransferase | 512 | 7.19 |
| **PA14_31750** | Unknown | Putative acyltransferase | 512 | 6.92 |
| **PA14_01670** | Unknown | Putative ATP-binding component of ABC transporter | 512 | 6.89 |

| | | | | |
|---|---|---|---|---|
| **PA14_49740** | Unknown | Uncharacterized protein | 222.990181 | 6.83 |
| **PA14_49480** | Unknown | ABC transporter domain-containing protein | 174.197333 | 6.78 |
| **PA14_45520** | Unknown | Putative plasmid partitioning protein | 259.177599 | 6.67 |
| **PA14_24590** | Unknown | Uncharacterized protein | 512 | 6.66 |
| **PA14_71910** | wbpZ | Glycosyltransferase WbpZ | 512 | 6.57 |
| **PA14_56070** | mvaT | Transcriptional regulator MvaT, P16 subunit | 15.482892 | 6.47 |
| **PA14_36420** | Unknown | Putative histidine kinase | 512 | 6.45 |
| **PA14_35050** | Unknown | Putative protease | 512 | 6.45 |
| **PA14_24650** | rmf | Ribosome modulation factor (RMF) | 512 | 6.43 |
| **PA14_44280** | rlmM | Ribosomal RNA large subunit methyltransferase M (EC 2.1.1.186) (23S rRNA (cytidine2498-2'-O)-methyltransferase) (23S rRNA 2'-O-ribose methyltransferase RlmM) | 256 | 6.43 |
| **PA14_00010** | dnaA | Chromosomal replication initiator protein DnaA | 171.196267 | 6.33 |
| **PA14_54770** | Unknown | Uncharacterized protein | 512 | 6.21 |
| **PA14_46930** | gltK | Putative permease of ABC transporter | 257.002852 | 6.16 |

| | | | | |
|---|---|---|---|---|
| **PA14_60650** | Unknown | Uncharacterized protein | 341.333333 | 6.13 |
| **PA14_53150** | Unknown | Probable ATP-binding/permease fusion ABC transporter | 512 | 6.05 |
| **PA14_24420** | Unknown | DUF58 domain-containing protein | 512 | 6.05 |
| **PA14_11730** | Unknown | Possible protein kinase | 256 | 5.79 |
| **PA14_18880** | nth | Endonuclease III (EC 4.2.99.18) (DNA-(apurinic or apyrimidinic site) lyase) | 512 | 5.77 |
| **PA14_59550** | Unknown | Uncharacterized protein | 512 | 5.74 |
| **PA14_35800** | Unknown | Uncharacterized protein | 512 | 5.69 |
| **PA14_49930** | Unknown | Uncharacterized protein | 256.5296 | 5.65 |
| **PA14_63530** | selB | Selenocysteine-specific elongation factor | 512 | 5.59 |
| **PA14_66650** | pilN | Type 4 fimbrial biogenesis protein PilN | 256.2648 | 5.48 |
| **PA14_29590** | Unknown | Putative transcriptional regulator | 8.90398237 | 5.47 |
| **PA14_25430** | lolC | Putative lipoprotein releasing system, permease protein | 172.078933 | 5.41 |
| **PA14_61790** | pth | Peptidyl-tRNA hydrolase (PTH) (EC 3.1.1.29) | 171.593248 | 5.39 |
| **PA14_42390** | exsA | Transcriptional regulator ExsA | 512 | 5.23 |
| **PA14_30260** | bpt | Aspartate/glutamate leucyltransferase (EC 2.3.2.29) | 512 | 5.22 |

| | | | | |
|---|---|---|---|---|
| **PA14_14830** | rlmN | Dual-specificity RNA methyltransferase RlmN (EC 2.1.1.192) (23S rRNA (adenine(2503)-C(2))-methyltransferase) (23S rRNA m2A2503 methyltransferase) (Ribosomal RNA large subunit methyltransferase N) (tRNA (adenine(37)-C(2))-methyltransferase) (tRNA m2A37 methyltransferase) | 512 | 5.21 |
| **PA14_13680** | Unknown | Putative short-chain dehydrogenase | 512 | 5.18 |
| **PA14_40840** | sohB | Putative protease | 14.6310844 | 5.11 |
| **PA14_58060** | Unknown | UPF0307 protein PA14_58060 | 512 | 5.09 |
| **PA14_11970** | Unknown | Putative 3-methyladenine DNA glycosylase (EC 3.2.2.-) | 512 | 5.07 |
| **PA14_08050** | Unknown | Putative tail fiber protein | 512 | 5.01 |

#### Table S3a: Fluorescence foci summary

| | DAPI | | | mApple | | |
|---|---|---|---|---|---|---|
| Strain | 1-foci | 2-foci | >2-foci | 1-foci | 2-foci | >2-foci |
| WT | 17±2 | 64±2 | 19±4 | - | - | - |
| *algP-mApple* | 14±1 | 55±2 | 32±3 | 5±3 | 53±2 | 41±4 |
| *mApple-algP* | 72±5 | 27±4 | 2±0.5 | 75±4 | 24±4 | 2±0.3 |
| *ΔalgP* | 77±0.2 | 22±0.2 | 1±0.3 | - | - | - |
| *algPΔCTD* | 71±4 | 28±4 | 2±1 | - | - | - |
| *ΔalgP* $P_{algP}$*:algP* | 33±4 | 62±3 | 6±2 | - | - | - |

#### Table S3b: Transmission Electron Microscopy Summary Data

| | WT | | ΔpolyP | |
|---|---|---|---|---|
| | 1.5h | 3h | 1.5h | 3h |
| Granule #/cell | 4.3±2.1 | 2.8±1.1 | 1.3±1.0 | 1.6±0.9 |
| Total granular volume/cell ($x10^{-3}$ **$\mu m^3$**) | 8.3±3.8 | 14±5.3 | 6.1±5.1 | 17±8.2 |
| Volume of largest granule/cell ($x10^{-3}$ **$\mu m^3$**) | 3.9±2.7 | 7.4±2.7 | 5.1±4.7 | 15±7.7 |
| Average granule volume (**$\mu m^3$**) ($x10^{-3}$ **$\mu m^3$**) | 1.9±1.9 | 4.8±3.2 | 4.0±4.4 | 10±9.0 |

#### Table S3c: Cell cycle exit

| | % >1 origin/cell | | % >0 fork/cell | |
|---|---|---|---|---|
| | 0h | 6h | 0h | 6h |
| WT | 86±2 | 10±6 | 68±18 | 3±2 |
| ΔpolyP | 88±5 | 87±4 | 74±5 | 35±3 |
| *ΔalgP* | 56±4 | 13±5 | 70±8 | 2±2 |

**Table S4a: Strains**

| Name | Genotype | Source |
|---|---|---|
| LR31; DKN263 | *P. aeruginosa* UCBPP-PA14 | |
| DKN303 | *E. coli* DH5α, pMQ30 | (2) |
| DKN1297 | *E. coli* DH5α (F− **Δ**(argF-lac)169 **Φ**80dlacZ58(**Δ**M15) glnV44(AS) **λ**− rfbC1 gyrA96(NalR) recA1 endA1 spoT1 thi-1 hsdR17 deoR), pUC18R6K-mini-Tn7T-Gm | (3) |
| DKN1298 | SM10, pTNS1 | (3) |
| DKN1299 | HB101 (F− **λ**− **Δ**(gpt-proA)62 leuB6 glnV44(AS) araC14 galK2(Oc) lacY1 **Δ**(mcrC-mrr) rpsL20(StrR) xylA5 mtl-1 recA13 hsdS20), pRK2013 pRK2013 has a ColE1 replicon and carries the RK2 tra genes and Tn903 (which is KanR) | (3) |
| LR135; DKN1729 | *P. aeruginosa* UCBPP-PA14 Δ*ppk1* Δ*ppk2B* Δ*ppk2C*; deletion of PA14_69230, PA14_33240, and PA14_19410 in DKN263 | (4) |
| LR79; DKN1730 | *P. aeruginosa* UCBPP-PA14 Δ*ppk2A* Δ*ppk2B* Δ*ppk2C*; deletion of PA14_01730, PA14_33240, and PA14_19410 in DKN263 | (4) |
| LR119; DKN1731 | *P. aeruginosa* UCBPP-PA14 Δ*ppk1* Δ*ppk2A* Δ*ppk2B* Δ*ppk2C*; deletion of PA14_69230, PA14_01730, PA14_33240, and PA14_19410 in DKN263 | (4) |
| LR177; DKN1736 | *P. aeruginosa* UCBPP-PA14 Δ*ppk1* Δ*ppk2A* Δ*ppk2B* Δ*ppk2C* attTn7:: mini-Tn7T-$Gm^R$ $P_{ssb}$*:ssb-mCherry* | (4) |
| LR229; DKN1762 | *E. coli* TOP10(F– *mcr*A Δ(*mrr-hsd*RMS-*mcr*BC) Φ80*lac*ZΔM15 Δ*lac*X74 *rec*A1 *ara*D139 Δ(*ara leu*) 7697 *gal*U *gal*K *rps*L (StrR) *end*A1 *nup*G), pLREX62 | (4) |
| LR322 | *E. coli* TOP10(F– *mcr*A Δ(*mrr-hsd*RMS-*mcr*BC) Φ80*lac*ZΔM15 Δ*lac*X74 *rec*A1 *ara*D139 Δ(*ara leu*) 7697 *gal*U *gal*K *rps*L (StrR) *end*A1 *nup*G), pLREX79 | This study |
| LR457 | *E. coli* TurboCells$^{TM}$ (F- recA1 endA1 hsdR17 supE44 thi-1 gyrA96 relA1 φ80lacZΔM15 Δ(lacZYAargF) U169), pLREX120 | This study |
| LR458 | *E. coli* TurboCells$^{TM}$ (F- recA1 endA1 hsdR17 supE44 thi-1 gyrA96 relA1 φ80lacZΔM15 Δ(lacZYAargF) U169), pLREX121 | This study |

| | | |
|---|---|---|
| LR461 | *E. coli* TurboCells$^{TM}$ (F- recA1 endA1 hsdR17 supE44 thi-1 gyrA96 relA1 φ80lacZΔM15 Δ(lacZYAargF) U169), pLREX124 | This study |
| LR462 | *E. coli* TurboCells$^{TM}$ (F- recA1 endA1 hsdR17 supE44 thi-1 gyrA96 relA1 φ80lacZΔM15 Δ(lacZYAargF) U169), pLREX125 | This study |
| LR467 | *P. aeruginosa* UCBPP-PA14 Δ*algP;* deletion of PA14_69370 | This study |
| LR469 | *P. aeruginosa* UCBPP-PA14 *algP*Δ*CTD;* truncation of 195 c-terminal amino acids (157-352) of AlgP (PA14_69370) | This study |
| LR471 | *P. aeruginosa* UCBPP-PA14 *algP::mApple-algP*; replacement of AlgP with chimeric fusion mApple-AlgP | This study |
| LR477 | *P. aeruginosa* UCBPP-PA14 Δ*ppk1* Δ*ppk2A* Δ*ppk2B* Δ*ppk2C* algP::algP-mApple; deletion of PA14_69230, PA14_01730, PA14_33240, and PA14_19410, replacement of algP (PA14_69370) with chimeric fusion algP-mApple | This study |
| LR491 | *P. aeruginosa* UCBPP-PA14 *algP::algP-mApple* ppk2A::ppk2A-mNeonGreen; replacement of AlgP (PA14_69370) with chimeric fusion AlgP-mApple, replacement of Ppk2A (PA14_01730) with chimeric fusion Ppk2A-mNeonGreen | This study |
| LR498 | *P. aeruginosa* UCBPP-PA14 Δ*ppk2a* Δ*ppk2b* Δ*ppk2c algP::algP-mApple*; deletion of PA14_01730, PA14_33240, and PA14_19410, replacement of AlgP (PA14_69370) with chimeric fusion AlgP-mApple | This study |
| LR500 | *P. aeruginosa* UCBPP-PA14 Δ*algP* (PA14_69370) attTn7:: mini-Tn7T-Gm$^{R}$ ParS$^{pMT1}$ $P_{ssb}$*:ssb-mCherry gfp-parB*$^{pMT}$*;* | This study |
| LR501 | *P. aeruginosa* UCBPP-PA14 Δ*algP* (PA14_69370) attTn7:: mini-Tn7T-Gm$^{R}$ $P_{algP}$*:algP;*algP complementation | This study |
| LR502 | *P. aeruginosa* UCBPP-PA14 *algP::algP-mApple*; replacement of *algP* with chimeric fusion *algP-mApple* | This study |
| LR503 | *E. coli* TurboCells$^{TM}$ (F- recA1 endA1 hsdR17 supE44 thi-1 gyrA96 relA1 φ80lacZΔM15 Δ(lacZYAargF) U169), pLREX132 | This study |
| LR504 | *P. aeruginosa* UCBPP-PA14 Δ*ppk1* Δ*ppk2B* Δ*ppk2C algP::algP-mApple*; deletion of PA14_69230, PA14_33240, and PA14_19410, | This study |

| |
|---|
| replacement of *algP* (PA14_69370) with chimeric fusion *algP-mApple* |

**Table S4b: Plasmids**

| Name | Genotype/Purpose | | Source |
|---|---|---|---|
| pMQ30 | Suicide vector | $Gm^R$ | (2) |
| pUC18R6K-mini-Tn7T-Gm | Mobilizable mini-Tn7 base vector with MCS | $Gm^R$ | (3) |
| PLREX9 | *ppk2a(*PA14_01730)*::ppk2a-mCherry*; pMQ30 derivative | | |
| pLREX62 | pUC18T-mini-Tn7T-$G^R$ $ParS^{pMT1}$ $P_{ssb}$ *SSB-mCherry GFP-ParB* $^{pMT1}$*;* pUC18R6K-mini-Tn7T-Gm derivative | $Gm^R$ | (4) |
| pLREX79 | *ppk2a(*PA14_01730)*::ppk2a-20aa-mNeonGreen*; pMQ30 derivative | $Gm^R$ | This study |
| pLREX120 | *algP* (PA14_69370) deletion vector; pMQ30 derivative | $Gm^R$ | This study |
| pLREX121 | *algP::algPΔCTD*; pMQ30 derivative, truncation of 195 c-terminal amino acids (157-352) of algP (PA14_69370) | $Gm^R$ | This study |
| pLREX124 | *algP(*PA14_69370)*::mApple-algP*; pMQ30 derivative | $Gm^R$ | This study |
| pLREX125 | *algP(*PA14_69370)*::algP-mApple*; pMQ30 derivative | $Gm^R$ | This study |
| pLREX132 | pUC18T-mini-Tn7T-$G^R$ $P_{algP}$*:algP; algP* complementation, pUC18R6K-mini-Tn7T-Gm derivative | $Gm^R$ | This study |

**Table S4c: Primers**

| Name | Purpose | Sequence |
|---|---|---|
| LRPR894F | Construction of *algP* derivatives at native locus | GACCATGATTACGAATTCGAGCTCGGTACCTCAGCGGACGCCCAGCAGGTCGATCTCG |
| LRPR912R | Construction of **pLREX120** [Δ*algP*] | CGCAGCCGGCTTGGCCGCAGGCTTCATGACGTGCCTCCAGGCGGACGTGGTTGCGCCC |
| LRPR909F | Construction of **pLREX120** [Δ*algP*] | GGCGCAACCACGTCCGCCTGGAGGCACGTCCTAAGGCGCTGTCTGCAAAGCCGCCGAGCC |
| LRPR899R | Construction of *algP* constructs at native locus | CGTTGTAAAACGACGGCCAGTGCCAAGCTTATGACTGGAAATGTCTGGAAATTCGCGGTG |

| | | |
|---|---|---|
| LRPR933R | Construction of **pLREX121** [*algP∆CTD*] | GGCTCGGCGGCTTTGCAGACAGCGCCTTAGTTACGCCGCTACGGTTTTCATCGCAGGC |
| LRPR932F | Construction of **pLREX121** [*algP∆CTD*] | GCCAAGCCTGCGATGAAAACCGTAGCGGCGTAACTAAGGCGCTGTCTGCAAAGCCGCCG |
| LRPR904R | Construction of **pLREX124** [*algP::mApple-algP*] | GGCCATATTGTTTTCCTCGCCCTTCGACACCATGACGTGCCTCCAGGCGGACGTGGTTGC |
| LRPR905F | Construction of p **pLREX124** [*algP::mApple-algP*] | GCAACCACGTCCGCCTGGAGGCACGTCATGGTGTCGAAGGGCGAGGAAAACAATATGGCC |
| LRPR906R | Construction of **pLREX124** [*algP::mApple-algP*] | GGTGCAAGGGGGTGGTGACGGGCTTCTTGTTGGCCGATTTATAGAGTTCGTCCATCCCC |
| LRPR907R | Construction of **pLREX124** [*algP::mApple-algP*] | GGGGATGGACGAACTCTATAAATCGGCCAACAAGAAGCCCGTCACCACCCCCTTGCACC |
| LRPR901R | Construction of **pLREX125** [*algP::algP-mApple*] | GAACTCTTTGATGATGGCCATATTGTTTTCCTCGCCCTTCGACACGGAGGCGCTGGTCGGGGCGGCGCCGTTGCTGCCCG |
| LRPR900F | Construction of **pLREX125** [*algP::algP-mApple*] | GCAACGGCGCCGCCCCGACCAGCGCCTCCGTGTCGAAGGGCGAGGAAAACAATATGGCC |
| LRPR903R | Construction of **pLREX125** [*algP::algP-mApple*] | GCTTGGCTCGGCGGCTTTGCAGACAGCGCCTTAGTTATTTATAGAGTTCGTCCATCCCC |
| LRPR898F | Construction of **pLREX125** [*algP::algP-mApple*] | GACGTGATGGGTATGGATGAACTCTATAAGTCGGCCAACAAGAAGCCCGTCACCACCC |
| LRPR956F | Construction of **pLREX132** [*$P_{algP}$:algP]* | CTTATCTGGTTGGCCTGCAAGGCCTTCGCGAGGTACCGGGGCGGTTTCTCCAGACGAATC |
| LRPR955R | Construction of **pLREX132** [*$P_{algP}$:algP]* | GTGGATCCCCCGGGCTGCAGGAATTCCTCGAGAAGCTTGGGTTAGGAGGCGCTGGTCGGG |
